# Reduced Telomerase Interaction with Telomeres Alters Meiotic Chromosome Motion and Gamete Viability

**DOI:** 10.1101/654376

**Authors:** Dana L. Smith, Ashwini Oke, Michael Pollard, Carol M. Anderson, Tangna Zhuge, Phoebe Yam, Tatiana Gromova, Kaylynn Conant, Daniel B. Chu, Neem J. Patel, Fernanda Gonzalez, Caitlin Stoddard, Sean Burgess, Andreas Hochwagen, Wallace F. Marshall, Elizabeth Blackburn, Jennifer C. Fung

## Abstract

We report a role for telomerase, beyond its known function of telomeric DNA end extension, in maintaining normal chromosome dynamics during meiosis in *Saccharomyces cerevisiae.* When telomerase at telomeres was reduced by various genetic means, increased frequencies of crossover and noncrossover recombination events occurred. To investigate the mechanism of this increased meiotic recombination, we examined the kinetics of meiosis events, and tracked the movement of chromosomes in live cells during meiotic prophase. Cytoskeletal forces acting on telomeres during meiosis have been shown to promote active chromosome motion needed to pair homologous chromosomes. Here we show that changes in telomerase interaction with telomeres using a tlc1-11 mutant result in altered meiotic motion. Specifically, reduction in telomerase at telomeres leads to a decreased frequency of high velocity chromosome pulls. In the tlc1-11 mutant, we see earlier synapsis and increased genome-wide recombination for the majority of the cells and lower gamete viability. Notably, homologous pairing is not delayed unlike other telomere binding mutants. Although synapsis initiates earlier, the overall timing of synapsis remains the same, except for a subset of cells that do not exit meiosis I. Together, these results suggest that the strong pulling component of the active chromosome motion promotes homolog pairing fidelity, likely by pulling apart improperly associated regions. Our combined observations are consistent with a model in which telomerase-mediated telomeric anchoring to the nuclear envelope helps engage and properly transmit cytoskeletal forces to chromosomes. Thus, telomerase contributes to efficient chromosome movements leading to normal gamete viability.

## Introduction

Segregation of homologous chromosomes is a key step in meiosis in which diploid cells halve their genome to produce haploid gametes. In humans, a leading cause of infertility, miscarriages and developmental disabilities is the failure to correctly segregate homologous chromosomes, resulting in aneuploidy (1). An essential event for correct chromosome segregation is the precise pairing of homologs prior to crossover recombination. Given the large number of homologous sequence tracts interspersed genome-wide among non-cognate chromosomes, an intriguing question is how homologs reliably find their correct partners (2).

Many organisms show some degree of premeiotic homolog associations that are disrupted through unknown mechanisms prior to stable meiotic pairing (3). This phase is followed by the accumulation of chromosomal associations by recombination-independent means (e.g. pairing centers, centromere coupling, bouquet) (4–7). Once chromosomes are brought together in close proximity, homology assessment ensues. In most organisms, homology assessment is recombination dependent (*C. elegans* and *Drosophila* being notable exceptions). Recombination initiates with Spo11-induced breaks (8) that are resected to create 3’ single-stranded DNA tails. These DNA tails invade another chromosome to assess homology while also initiating DNA repair to form noncrossovers (NCOs) or crossovers (COs) between homologs that eventually form chiasmata.

Telomeres play an important role in chromosome reorganization via their engagement with the nuclear envelope (NE) during meiosis (9, 10). During meiotic leptotene, telomeres cluster in proximity to the centrosome forming the telomere bouquet (11). Although the bouquet disperses after zygotene, telomere-mediated connections to the SUN/KASH domain proteins in the NE remain throughout meiotic prophase, connecting chromosomes with the cytoskeleton to promote chromosome motion (12) Initially observed in fission yeast (13), then in budding yeast, telomere-led chromosome motion has been reported in other species including plants, flies, worms and mammals. In budding yeast, chromosomes move on average ∼ 0.3 μm/sec but sporadically move at higher velocity (>1 μm/sec) in telomere-led pulls (14, 15). The distinct kinds of motion depends on whether or not a chromosome engages with the cytoskeleton via its telomeres. Many possible roles for the overall motion have been proposed (Figure 1A). All the necessary components that allow this force to be applied to telomeres are still unclear, as are the roles of this motion and whether motion is needed primarily for increasing collisions of homologous regions, testing of pairing fidelity, resolving interlocks or some combination of each role. Motion was first assumed to expedite contact between homologous regions (13), but current models propose that these forces may act to limit incorrect associations as well (16–18). Experiments perturbing the pulling component of the motion predicts distinct consequences for meiosis depending on its functional role(s) (Figure 1B).

**Figure 1.**
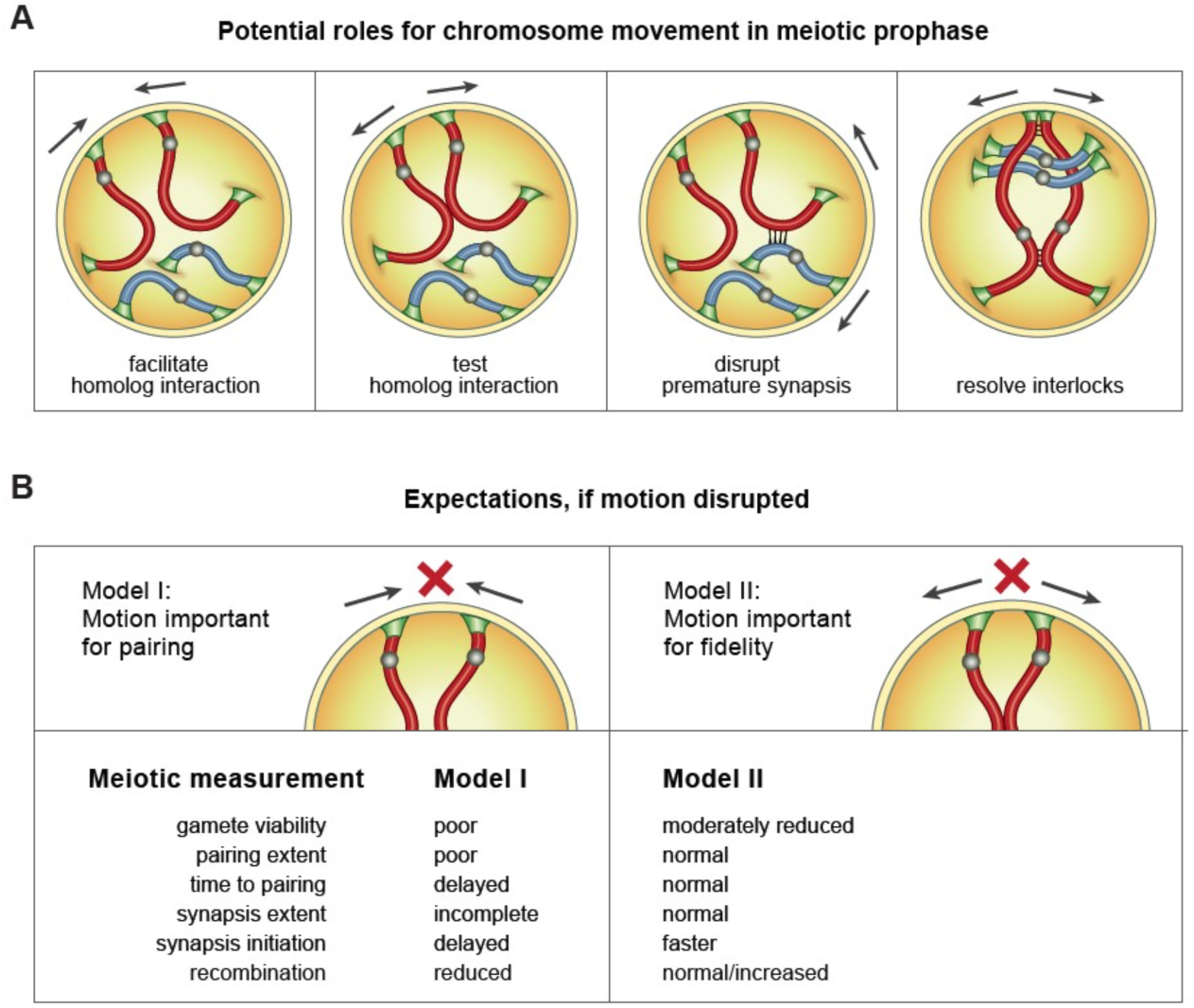
Models for role of active motion in meiosis. **A)** Four models for the role of telomere-led chromosome movement: 1) motion brings homologs in close proximity; 2) motion pulls chromosomes apart to test homolog fidelity; 3) motion disassociates inappropriate synapsis and 4) motion removes interlocks or entanglements. **B)** Different simple expectations for models I vs. II upon perturbation of motion for the majority of chromosomes.

In mitotic budding yeast cells, telomeres are tethered to the NE via two complexes (19) (Figure 2A). One complex tethers the subtelomeric chromatin and is made up of Sir4, a chromatin silencing protein, and Esc1, a protein found exclusively at the inner face of the NE (20). A second complex includes components of the telomerase holoenzyme (Figure 2A) and anchors telomere distal ends to the NE. This telomerase-dependent attachment relies on the interactions of yKu with TLC1 (the RNA component of telomerase) and of Est1 (a core protein component of telomerase) with the Mps3 SUN domain protein (19). Ndj1 is an additional component of the second complex that only plays a part during meiosis. Though important in mitosis for telomere transcription and maintenance (21), the functional importance of these first two complexes have yet to be fully explored during meiosis.

**Figure 2.**
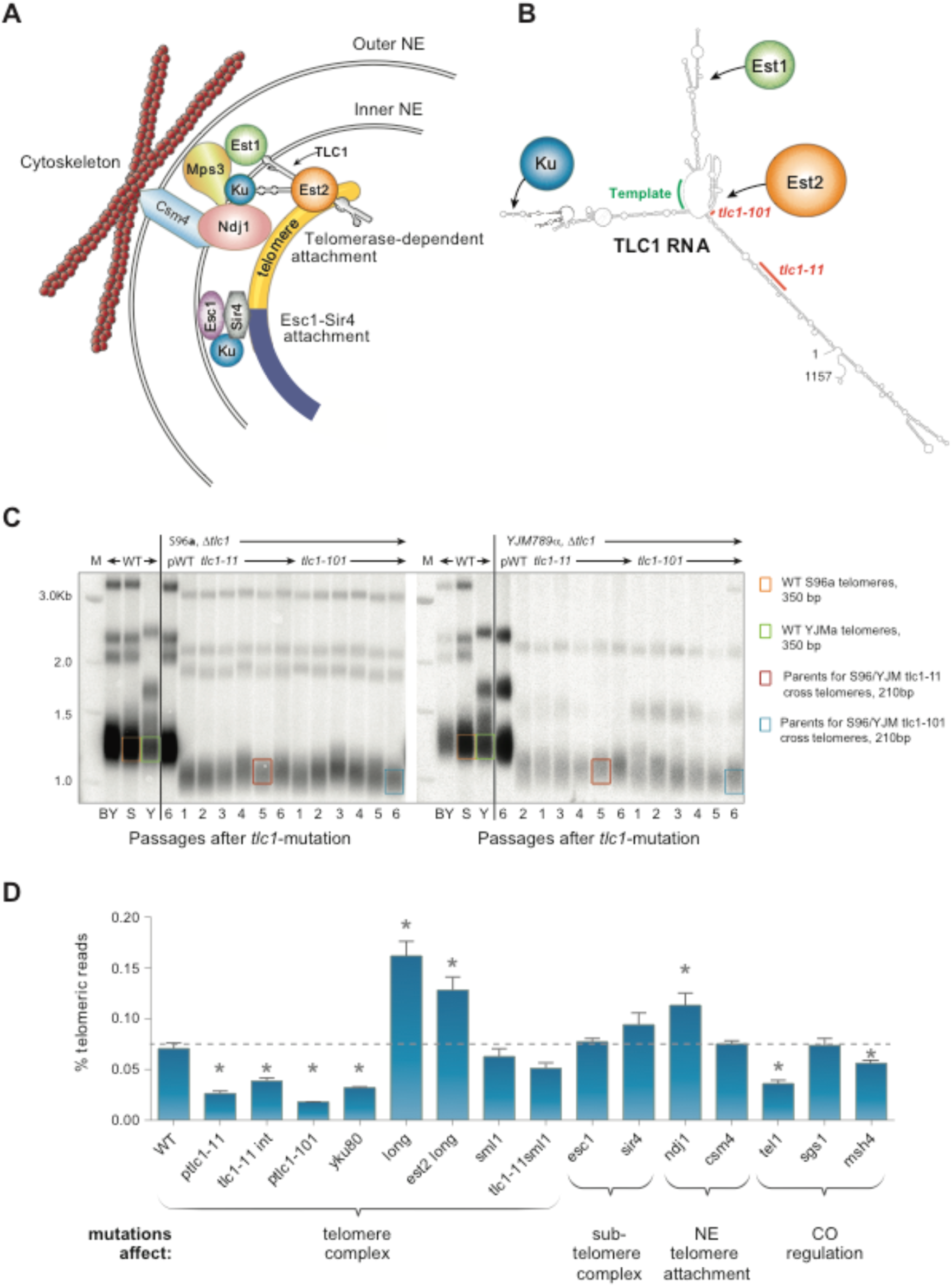
Mutations in telomerase and telomere maintenance components used to distinguish effects of altering two types of telomere attachments. **A)** A model for nuclear envelope (NE) telomere anchoring in mitotic cells, with speculative inclusion of meiotically-expressed Ndj1. Two types of telomere attachment to the NE are known: telomerase-dependent and subtelomeric. The telomeric tip is attached through components of the telomerase holoenzyme whereas Esc1 and Sir4 facilitate attachment of the subtelomere. TLC1, the RNA portion of telomerase, binds both Ku and Est1; Est1 binds Mps3, a protein firmly anchored in the NE. Ndj1 acts at the telomerase-dependent attachment during meiosis via interaction with Csm4. **B)** Model showing the secondary structure of the TLC1 RNA with binding sites for Ku, Est1 and Est2. (Red) *tlc1-101* and *tlc1-11* mutations. **C)** Southern blot analysis of terminal XhoI restriction fragments to calculate telomere size. Generation of mutant alleles of *TLC1*. Cells were serially passed 6 times (∼120 generations). S96a and YJM789 *tlc1* mutants with similarly short telomeres were chosen for mating and subsequent sequencing of gametes. DNA was probed with an γ32P-labeled 5’-(TGTGGG)4-3’ Y’-specific probe. The lowest species represents the DNA fragment containing the terminal telomeric repeats. WT telomeres are about 350 bps; *tlc1-11* and *tlc1-101* are 210 and 160 bps respectively (For long telomere mutants see Figure S1). **D)** Relative telomere lengths determined from whole genome sequencing of WT and mutant spores in S96/YJM789. Yeast telomeres reads consist of a degenerate repeating sequence, T followed by 1-3 Gs, TG-rich and AC-rich (complementary strand). Non-telomeric reads with long single letter runs arising from sequencing artifacts were removed.

To investigate the role of telomerase at telomeres in meiotic prophase motion, we deleted telomerase or genetically manipulated it to reduce the levels of telomerase at telomeres thereby modulating the engagement of telomeres to the NE. Here we present evidence supporting a model in which telomerase-dependent engagement is important for meiosis and gamete viability. Specifically, partial reduction or complete loss of telomerase at telomeres in meiotic prophase resulted in fewer and less effective interactions of the telomere with the cytoskeleton, greatly reducing the number of “pulls” a chromosome experiences and moderately decreasing the strength of the pulls. Reduction of telomerase also led to earlier synapsis, higher genome-wide recombination event frequencies and lower spore viability. Together, these findings suggest that cytoskeletal mediated chromosome mobility function not to bring homologous chromosomes together but rather favors alternative models (Figure 1B) (e.g. to test the fidelity of homologous associations by pulling apart incorrect associations

## Results

### Mutants that affect levels of telomerase at telomeres

To better understand the role of telomerase in NE attachment and motion in meiosis, we mutated two complexes known to anchor telomeres to the yeast NE in mitotic cells: the distal telomerase-associated complex containing core ribonucleoprotein components TLC1 and Est2 as well as the telomerase recruitment protein, yKu, and the sub-telomere-binding complex containing Sir4 and Esc1 (Figure 2A, Table S1).

While deletion of *SIR4* or *ESC1* still allows long-term viability, deleting telomerase holoenzyme genes causes telomeres to progressively shorten, resulting in eventual cell senescence after many cell divisions. Therefore, to study the effect on meiosis without this added complication, we first used two previously-characterized mutations in *TLC1, tlc1-11* and *tlc1-101,* that only partially reduce levels of assembled Est2-TLC1 complex (22) (Figure 2B, map adapted from Hass and Zapulla 2015 (23)). These two *tlc1* hypomorphic mutants are viable long-term and maintain stable but shorter telomeres, of average length 210 bp (*tlc1-11*) and 160 bp (*tlc1-101*), compared to the WT length of about 350 bp (Figure 2C). *tlc1-101* is a 3-nucleotide substitution of bases 814-816, ACU->UGA, in a region critical for the binding of *TLC1* RNA to Est2p, the enzymatic core protein component of telomerase. *tlc1-11*, a 30-bp deletion of nucleotides 850-880, reduces steady-state levels of TLC1. Both *tlc1* mutants result in less assembled TLC1-

### Est2 ribonucleoprotein (RNP) core complex, thereby predicting reduced telomerase holoenzyme at telomeres

Independently, to test complete deletion of telomerase function in meiosis without complications due to excessive telomere shortening, we also created strains with greatly pre-elongated telomeres. This was achieved by using the Cdc13-Est2 fusion protein, which lengthens telomeres by constitutively tethering telomerase to telomere ends (24). We first lengthened telomeres to five times their normal length, before removing the *CDC13-EST2*-expressing plasmid. Using this technique, we examined meiosis in strains with long telomeres, either with endogenous telomerase (*EST2)* “*long*” or without (*est2Δ)* “*est2 long*” (Figure S1A). Long telomeres alone have been shown to have reduced levels of telomerase (25, 26) so we expect to see similar phenotypes in both “*long*” and “*est2 long*”.

We constructed additional mutant strains previously known to affect telomere anchoring to the NE: *yku80Δ*, *ndj1Δ and csm4Δ.* Ndj1 is a meiosis-specific tether (27) required for telomere-NE attachment (28). Csm4 protein mediates the attachment to the cytoskeleton and is important for telomere clustering (15, 29, 30).

We verified the relative lengths and sequence patterns of telomeres in our strains using whole genome sequencing, by identifying and quantifying telomeric reads that exhibited at least a 95% threshold for TG composition (Figure 2D and S2B; see Methods). In all these mutants, telomere sequences showed no statistical differences in the patterns of the irregular *S. cerevisiae* telomeric DNA repeat unit (G1-3T) from wild type. This assay also confirmed the lengths of previously reported short telomere mutants (*tlc1-11, tlc1-101, tel11,*), normal length telomere mutants (*sir41* and *esc1Δ*) and longer telomere lengths in the *long* mutants described above (Figure S1B). *ndj1Δ* also exhibited longer telomeres, in line with Ndj1’s role in resetting telomere size by telomere rapid deletion (31). Deletion of Sgs1 has been reported to exacerbate telomere shortness of some mutants (32). Using this analysis, we show that *sgs1Δ* alone has normal telomere length but *msh4Δ* leads to shorter telomere length.

### Mutations disrupting telomerase-associated attachment reduce gamete viability

Although the *tlc1* strains we examined have normal mitotic cell growth (Lin et al., 2004), in meiosis, spore viability was reduced about 20% (Figure 3 and Table S2). Mutants with shorter telomeres (*tlc1-11* and *tlc1-101, yKu80Δ and tel1Δ*) as well as strains with long telomeres with or without Est2, are predicted to have reduced telomerase engagement at telomeres. For *sml1 tlc1-11*, the deletion of *sml1* (which increases dNTP levels and is likely to increase the polymerizing action of telomerase and hence engagement of telomerase at telomeres) produced a corresponding increase in spore viability compared with *tlc1-11* alone (33–35) (Figure 3 and Figure S2). In contrast, *sir4Δ* and *esc1Δ* showed no effect on spore viability in either the homozygous BR1919-8B (Figure 3) or hybrid S96/YJM789 strains (Figure S2). These findings suggest that optimally efficient completion of sporulation requires the components of the telomerase complex, but not the subtelomeric Sir4-Esc1 complex.

**Figure 3.**
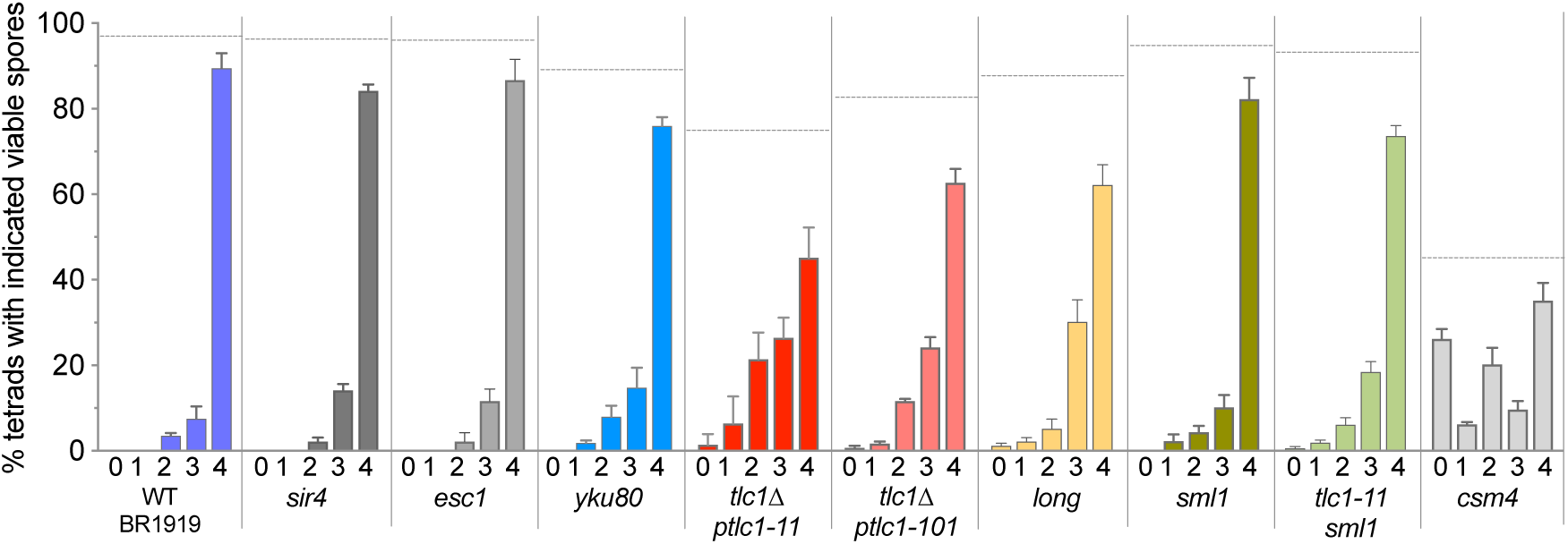
Analysis of spore viability of *tlc1-11*. Spore viability of telomeric binding protein mutants measured in strain BR1919-8B. Spore viability, % asci with 0,1,2,3 or 4 viable spores as indicated for each genotype below the histograms. Horizontal dotted line indicates total spore viability. (See Figure S3 for spore viability in SK1 background and Table S2 for summary of all mutants in each background.) n >200 tetrads per strain, averaging sets of 40 tetrads per plate; error bars represent SEM.

### Higher recombination caused by hypomorphic telomerase

Due to the reduction in gamete viability, we investigated whether recombination was affected in tlc1-11 and comparing it to several other mutants (see Figure 4), for altered recombination parameters. To gain an accurate representation of COs and NCOs genome-wide, we first measured recombination by whole genome sequencing (RecSeq) in the S96/YJM789 hybrid strain containing over 60,000 SNPs (45). Mutants that result in less telomerase at telomeres (whether or not telomeres were made short or elongated), as well as mutants in components that affect telomerase-dependent but not the sub-telomeric NE attachment, showed a genome-wide increase in COs (10-60%) (Figure 4A). NCOs also increased in proportion to COs (Figure 4B). The increased recombination was not restricted to telomeric regions but occurred chromosome-wide (Figure 4C). In contrast, *Δtel1* and *Δsgs1* mutations, while increasing COs, also altered the ratio of COs to NCOs, indicative of changes in CO regulation (46), suggesting that this aspect of CO regulation is unaffected in telomerase mutants. Interestingly, *esc1Δ* also resulted in a lower CO/NCO ratio, potentially suggesting a role for Esc1 in recombination. Contrary to the expectation of altered recombination, the distribution of recombination signatures was identical between *tlc1-11* and WT. (Figure 4D). Finally, in all the telomerase mutants, overall interference decreased moderately (Figure 4E). Together with the observation of early synapsis, the high level of error-free recombination suggests that perturbing the pulling forces on chromosomes, through diminished telomerase at telomeres, allows for the chromosomes to form stable associations earlier, permitting more interhomolog recombination to occur for the majority of the chromosomes. This is consistent with a model in which motion is needed to dissociate chromosomes as a means to test homology, whether they correctly or incorrectly pair.

**Figure 4.**
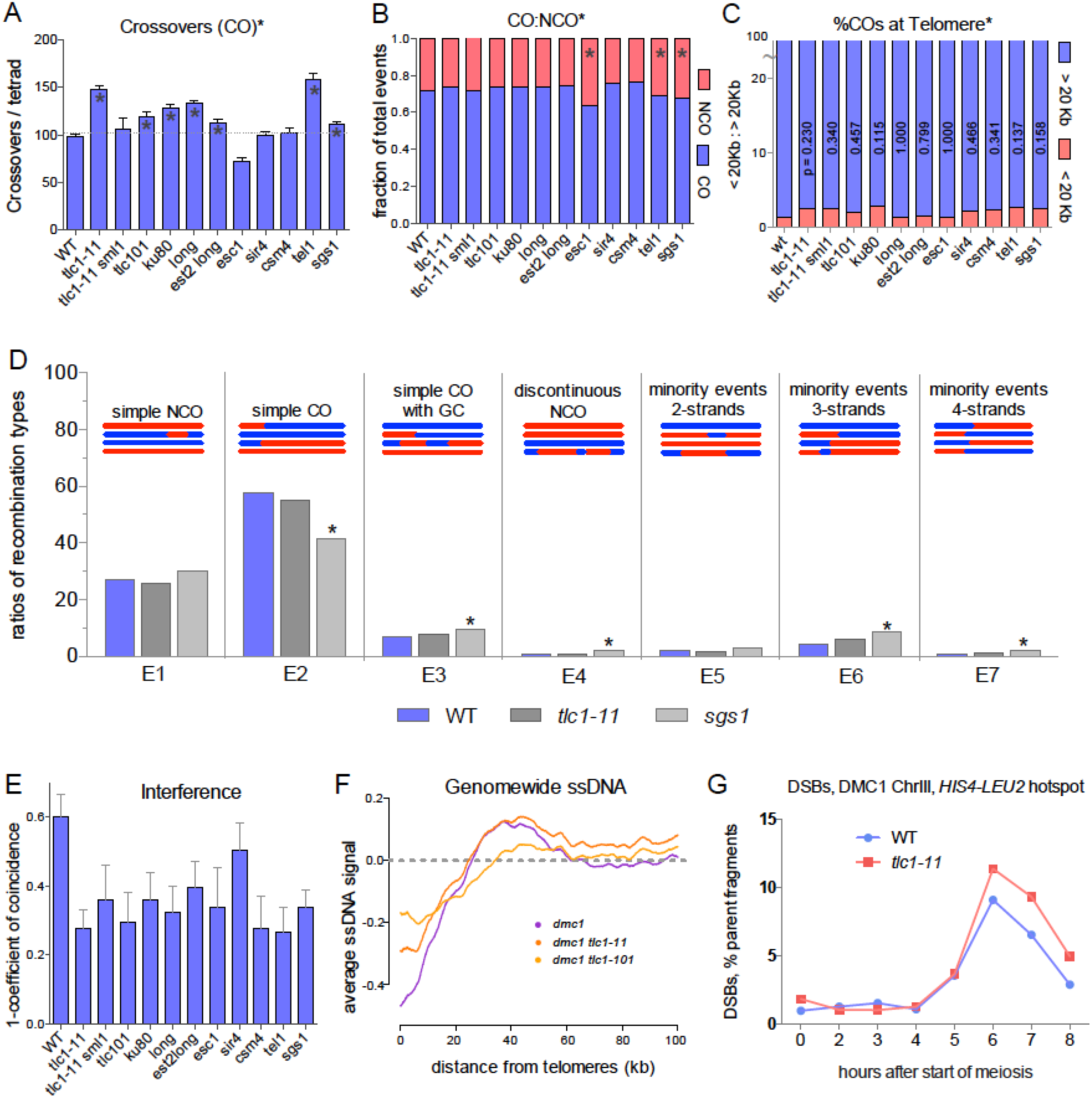
Higher recombination is observed for mutants with reduced telomerase. **A)** More COs in telomerase deficient mutants. Average COs per meiosis obtained from sequencing analysis. * chromosome 12 is removed from all strains for comparison purposes due to YJM789 chromosome 12 disomy in some strains. **B)** Ratio of total COs vs.NCOs (CO:NCO). **B)** CO frequency at telomere proximal vs. distal. P-values are from chi-square analysis of COs < 20Kb from the telomere end (red) and COs > 20Kb from the end (blue) for each mutant and WT. **C)** Comparison of recombination signatures between WT, *tlc1-11* and *sgs1***. E)** Average CO interference calculated from coincidence of adjacent intervals genome-wide. **F)** DSB levels measured using genome-wide ssDNA signal in *dmc1* SK1 as a function of distance from telomere. **G)** Timing and frequency of DSBs as percentage of parent fragments at the *HIS4-LEU2* DSB hotspot on chromosome III in a DMC1+ SK1 strain, (See Figure S6)

A CO increase can reflect either an increase in DSBs or a change of bias toward interhomolog repair of breaks. We measured DSBs by genome-wide detection of ssDNA in a synchronizable SK1 *dmc1Δ* background that allows ssDNA regions to be stably maintained at the sites of DSBs. When the DSBs of chromosome VIII were measured in *dmc1Δ*, neither *tlc1-11 dmc1Δ* nor *tlc1-101 dmc1Δ* showed more DSBs than *dmc1Δ* alone (Figure 4F, Figure S5A, B). Although deletion of *DMC1* facilitates the preservation and detection of DSBs, it can potentially mask DSBs introduced later in meiosis (47). We therefore measured DSBs at a single break site, the *HIS4*-*LEU2* hotspot on chromosome III, in SK1, without perturbing DSB processing (*DMC1*). Again, the number of DSBs in the *tlc1* mutants is similar to WT (Figure 4G, Figure S5C and D). We conclude that the increased recombination is due to a change in bias toward interhomolog DSB repair.

### Hypomorphic telomerase mutants result in major changes in the frequency and minor changes in the velocity of chromosome “pulls”

Having established that several mutants that have reduced telomerase presence have similar effects on spore viability and recombination in the final products of meiosis, we directly analyzed whether this affected chromosome mobility. We first investigated whether meiotic chromosome motion is disrupted in *tlc1-11* by imaging a long single chromosomes using Zip1-GFP in both wild type and *tlc1-11* 1919-8B strain backgrounds (Methods). Zip1 is a synaptonemal complex (SC) protein responsible for synapsing paired homologs along their lengths (36). In WT, the sixteen synapsing chromosomes in a compact nucleus make it difficult to definitively track the movements of a single chromosome. Thus, we imaged chromosome motion in a *zip3Δ* mutant, which less efficiently initiates synapsis such that although multiple chromosomes are synapsed, occasionally nuclei containing a single synapsed chromosome are observed (37–39)(Figure 5A), thereby allowing for clear analysis of chromosome motion (40)(Movies Figure S3A-B [zip3 TLC1.mov and zip3 tlc1-11.mov]). We determined the volume of the nucleus using the Zip1-GFP background signal from which the nuclear surface was calculated ((40)). To quantitatively measure any change in chromosome dynamics along the length of the synapsed chromosome, each chromosome was spatially digitized and then tracked through time using a semi-automatic method using intensity convolution and thresholding to compensate for optical distortion along the axis of microscopy (Figure 5B). Figure 5C shows that a given chromosome can traverse throughout the nucleus and is not confined to a limited region of the nucleus. For long chromosomes, the cumulative distance that telomeric regions (red and blue) travel is much greater than the cumulative distance of the midpoint (green), which remains predominantly in the center of the nucleus (Figure 5D). In all nuclei in which a chromosome was tracked, telomeric regions - whether from long or short chromosomes - never moved into the interior of the nucleus but remained on the nuclear surface. Moreover, each telomeric end of a chromosome moved independently of the other and motion is observed to be reduced further away from the telomere ends (Figure 5D), suggesting that the cytoskeletal forces at telomere ends are not well transmitted along the full length of long chromosomes. Consistent with earlier studies of meiotic chromosome motion (14, 15, 41), we found that chromosome motion in *zip3Δ TLC1* and *zip3Δ tlc1-11* is episodic consisting of periods of relative quiescence (“dwells”, shown in blue) alternating with sharp changes in velocity (“pulls”, shown in red) (Figure 5E, 5F) with average telomeric velocity of 0.3 μM/sec and pull velocities reaching > 1.0 μM/sec on par with velocities measured in these earlier studies. In Figure 5F, an example of a velocity profile over a short time interval is shown. We calculated the average kinetic parameters of chromosomal motion for each strain (Figure 5G) from all velocity profiles, defining a “pull” to be any peak greater than 10% of the maximum velocity observed (dotted line in Figure 5F). The “pulls” result from the chromosome end interacting with the cytoskeleton surrounding the NE (14). As summarized in Figure 4G, both the frequency and maximum velocity of the pulls are reduced in *tlc1-11 zip3Δ*, indicating that the mutation affecting telomerase-dependent attachment does perturb motion.

**Figure 5.**
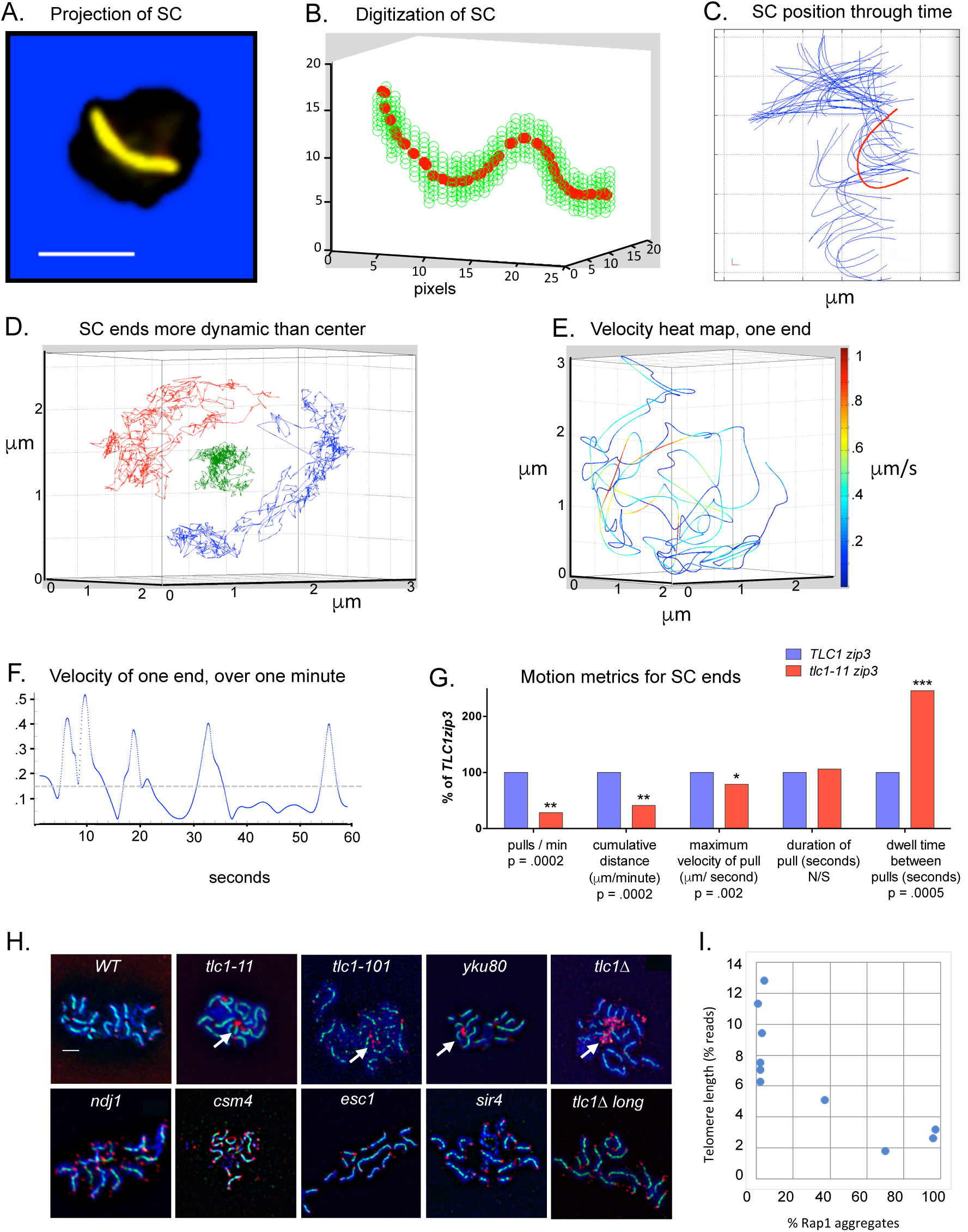
Analysis of telomere-led chromosome motion. **A)** Projection of a chromosome from a BR1919-8B *zip3* meiotic nucleus containing a Zip1-GFP fusion highlighting extent of synapsis for a single chromosome (Scale bar, 2μm). **B)** Digitized contour of the SC at one time point in 3D. Green circles represent segmented Zip1 intensity used to trace axial path of the SC through space (red circles) fitted to a quintic spline. **C)** Digitized contours throughout one time course. Red highlights a single contour. **D)** Ends move more than center of chromosome. Example from a *tlc1-11 zip3* strain. Chain of displacement vectors plotted for each end of a chromosome (red and blue) and for the mid-point (green). **E)** Velocity heat map. Complete path traversed by the one telomeric end depicted in red in D in which color is proportional to velocity in a six-minute timecourse. **F)** Changes of velocity of one end in one minute. Velocity of one end shown in E calculated from spline fit to particle path. Pulls are defined as periods of high velocity motion > 10% max velocity. Dwells are the periods between pulls. **G)** Changes of velocity magnitude and frequency in *tlc1-11*. Comparison of normalized SC motion metrics in *tlc1-11 zip3* (n=53 pulls over 29 minutes from 11 cells) or *zip3* (n=154 pulls over 24 minutes from 4 cells). * significant P-values (t-test). **H)** Immunostained spread pachytene chromosomes. DAPI (blue) DNA, Zip1-GFP (Green) and Rap1 (Red). White arrows indicate representative telomeric aggregates of Rap1. **I**) Rap1 aggregation vs telomere length.

### A telomerase hypomorph does not alter telomere end declustering

In budding yeast, telomeres cluster in leptotene to form the meiotic bouquet, which then disperses (declusters) after zygotene. The cumulative distance traveled for telomere ends in *tlc1-11* is less than for WT (Figure 5G), which not only indicates motion impairment but also raised the question of whether the telomeric ends decluster normally. To assay telomere declustering, we analyzed the frequencies at which two clustered (<0.3 μm apart) chromosome ends disperse during meiosis in live *zip3* vs. *zip3 tlc1-11* cells. In *tlc1-11* (n=24) and WT (n=38) nuclei, all chromosomes ends that were initially clustered could disperse at least 1 μm apart with similar timing, indicating that declustering was not impaired in the mutant. Initial experiments using Rap1 to visualize telomere clusters instead revealed that observed Rap1 aggregation is not necessarily related to the level of telomere clustering as typically suggested (42) but was inversely correlated to telomere length in the various mutants (Figure 5H, 5I,Figure S1B) This suggests that the formation of Rap1 clusters are the result of loss of telomere length rather than an indication of telomeric clustering.

### The hypomorphic telomerase mutant perturbs the quality of the engagement of chromosomal termini with the cytoskeleton during meiotic prophase I

We tested if the reduced frequency of telomere pulls was a consequence of perturbed telomere attachment to the NE and/or to the cytoskeleton. First, if *tlc1-11* telomerase caused NE detachment, as reported for *ndj1Δ* (28),telomeres should localize towards the interior of the nucleus as well as the periphery. Unlike *ndj1Δ*, chromosome ends in *tlc1-11 zip3Δ* remained only at the nuclear periphery (Figure 4D). Therefore, we tested whether *tlc1-11* affects the ability of the telomere to maintain attachment to the cytoskeleton, assessed by analyzing telomere motion parameters. Less efficient initial attachment events would cause less frequent (longer dwell times) but normal pull durations. Strikingly, *tlc1-11 zip3Δ* showed vastly longer dwell times between pulls (Figure 5G), indicating impaired ability to initiate pulls compared to WT. Earlier cytoskeletal detachment would result in shorter pull durations. Pull durations were the same with or without the *tlc1-11* (Figure 5G), suggesting that, once engaged, *tlc1-11* is able to maintain attachment to the cytoskeleton. However, the reduction of average maximal velocity to 75% of WT (Figure 5G) and less frequent pulls suggests that although attachment occurs, it is not fully functional in *tlc1-11*.

### Homolog pairing kinetics are not altered by mutations that reduce telomerase at telomeres

If chromosome pulls help homologs locate their partners more efficiently, disruption of the pulls may lead to slower pairing kinetics. Alternatively, if this chromosome pulling motion is mainly used to test homolog interaction and disrupt pairing mistakes, pairing kinetics should be less affected by reduced telomere motion especially if homology correction is a low frequency event. To distinguish between these possibilities, we examined the timing and extent of homologous pairing by fluorescently marking a site near the centromere of chromosome V of each homolog with a 256-LacO array in cells also expressing GFP-LacI repressor protein. We analyzed pairing time-courses of WT, *tlc1-11* and *tlc1-101* in the highly synchronous SK1 strain. The *tlc1-11* and *tlc1-101* mutants both exhibited WT pairing kinetics (Figure 6A and B). Pairing kinetics of a fluorescently marked telomere proximal site were also similar in *tlc1-11*, *tlc1-101* and WT strains (Figure S5A). The unperturbed timing and extent of homologous pairing in the *tlc1* strains suggests that the pulling motion does not primarily speed up pairing, but may contribute to other potential functions (Figure 1B).

**Figure 6.**
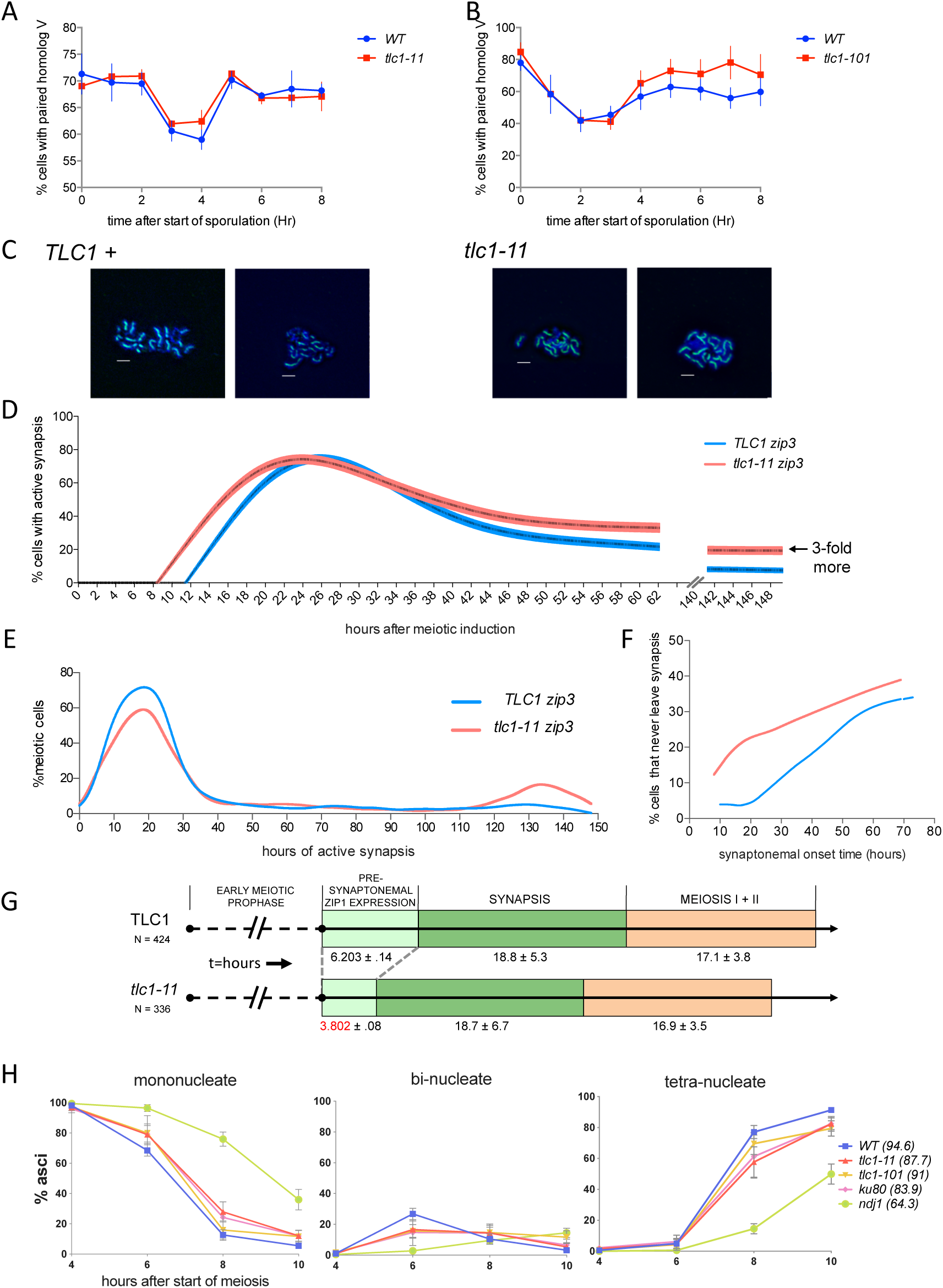
The *tlc1-11* mutant exhibits early synapsis but normal pairing. **A)** Time and extent of pairing for *tlc1-11*. Cells with LacO repeats inserted near the centromere of each homolog were fixed hourly after meiotic start. LacO foci detected with LacI-GFP are considered paired (1 spot) or unpaired (2 spots). Percentage of cells with paired homologs was determined for three strain crosses for each genotype n = 1000 cells/time point/ diploid strain. **B)** Time and extent of pairing for *tlc1-101*. Same as above. Scale bar = 2 μm**. C)** Extent of synapsis is normal in *tlc1-11*. Pachytene chromosomes were spread and immunostained against Zip1 for *WT and tlc1-11.* **D)** Synapsis initiates earlier in *tlc1-11*. Individual *tlc1-11 zip3* (n = 336) or *zip3* (n = 424) cells containing Zip1-GFP are tracked in vivo throughout the entire course of meiosis. Time to synapsis initiation was recorded. At the end of the live cell tracking a three-fold increase in cells were seen that still were in active synapsis. **E**) Analysis of synapsis duration. The time from synapsis initiation to synapsis termination was determined for *zip3* and *tlc1-11 zip3*. For *tlc1-11*, ∼20% of the population showed extremely long synapsis durations. **F)** Cells showing long synapsis durations start synapsis throughout time course, not only at late times. Time from induction to synapsis onset are recorded for cells with long synapsis durations. **G)** Early synapsis is due to shorter presynapsis period. t = hours. Summary of major periods of meiosis determined from single cell *in vivo* tracking until completion of meiosis. **H)** Meiotic progression of *tlc1-11, tlc1-101* and y*ku80*Δ and *ndj1Δ*. Frequency of nuclear division measured by DAPI in SK1 (n >3 x100cells/timepoint, error is SEM). (), indicates overall sporulation frequency.

### Hypomorphic telomerase results in precocious synapsis and failure to exit synapsis

To further test if telomeric pulling acts to physically disassociate incorrectly paired chromosomes apart, we next analyzed the timing of synapsis in *tlc1-11*. We predict that reduction of the telomeric pulling motion in *tlc1-11* should lead to earlier synapsis, since the time spent undoing incorrect associations and re-searching for correct homologs would be eliminated and synapsis could start earlier, even between nonhomologs. In pachytene chromosome spreads, *tlc1* mutants achieved full WT synapsis (Figure 6C). Furthermore, while synapsis extent was not perturbed, live imaging of individual synapsed pairs in single cells revealed that *tlc1-11 zip3Δ* started synapsis fully 2.5 +/-0.11 hours before *zip3Δ* (Figure 5D).

Despite initiating synapsis earlier, for the majority of individual *tlc1-11 zip3Δ* cells the absolute duration of synapsis was the same as in *zip3*Δ (18.7 vs. 18.8 hours) (Figure 6E). However, notably, ∼20% of *tlc1-11 zip3Δ* cells remained indefinitely in the process of synapsis assembly and disassembly, thus failing to exit meiosis I (Figure 5E). Figure 6F shows that the subpopulation that fails to exit synapsis includes cells that begin synapsis early as well as throughout the time course and thus is not the result of cells that enter late into meiosis and do not have time to finish. The inability to finish synapsis in this subset of the population is consistent with these cells containing unsynapsed chromosomes that prohibit meiotic progression, potentially through activation of the pachytene checkpoint (reviewed in (43)). Corroborating evidence for this interpretation was found in cell population meiotic progression assays done by quantifying percentages of cells at different stages at fixed time intervals throughout meiosis (Figure 6H). As a positive control, *ndj1Δ* showed a long delay in entering meiosis I, as expected (27, 44). The *tlc1-11*, *tlc1-101* and *ku80Δ* mutant cell populations each appeared to leave the one-nucleus stage to enter meiosis I only slightly later than WT; we note that this apparent “delay” in the average timing of progression is the observation expected in a cell population assay when a small subpopulation of cells fails to progress into meiosis I.

Consistent with model II in Fig S1B, precocious synapsis and the subpopulation exhibiting a delay or inability to enter MI is an expected outcome when the pulling motion is perturbed and mispaired regions are harder to disengage. Taken together, our results suggest that chromosome motion plays additional roles other than increasing active diffusion to enhance homology search.

### Pairing simulations predict that lowering the frequency of cytoskeletal attachments will cause increased association of chromosomes

Although *tlc1-11* alters telomere motion, it does not eliminate it as seen in the live movies (Figure S3B), so we wished to investigate whether the quantitative reduction observed in the frequency or velocity could in principle be sufficient to alter chromosome associations. We have previously used a computational model based on Brownian dynamics simulations (18) to show that forces applied to telomeres can, in principle, promote pairing fidelity by pulling weakly paired regions apart (48). By allowing chromosomes to associate reversibly (Figure 7A, movies Figure S6A-B), with the affinity of two chromosomes described by the rate at which paired loci become unpaired, we can ask what happens if we reduce motion. In simulations in which chromosomes are allowed to either pair with their correct homolog or an incorrect homeologous chromosome, with different off-rates, we frequently see that chromosomes will pair with the wrong chromosome even when the off rate is different by an order of magnitude (Figure 7B). Simulations showed that stretches of incorrect pairing were eventually pulled apart (Figure 7C), whereas in the absence of telomere forces, incorrectly paired regions remained associated for long periods of time (Figure 7D). The apparently stable association of nonhomologous regions, despite the high unpairing rate of individual loci, is due to avidity effects. When simulating homolog pairing with the observed reduction in the frequency of high velocity pulls, we found that chromosomes mis-associated 25% more frequently. Because our measurements showed not only a change in frequency but also a change in velocity, we also simulated homolog pairing with normal frequency of pulls but with a 75% reduction in velocity. This was also sufficient to increase the rate of mispairing. Reduction in either the frequency or velocity of the telomere pulling motions to an extent matching our experimental observations led to an increase in incorrect chromosome associations. Combining a 25% reduction in frequency with a 75% reduction in velocity, which mimics our experimental observations of the *tlc1* mutant in living yeast cells, led to a more dramatic effect (Figure 7E). We conclude that, based on our computational model, a quantitative reduction in either the frequency or velocity of telomere pulls, both of which we observed in *tlc1-11* mutants, can increase mispairing rates.

**Figure 7.**
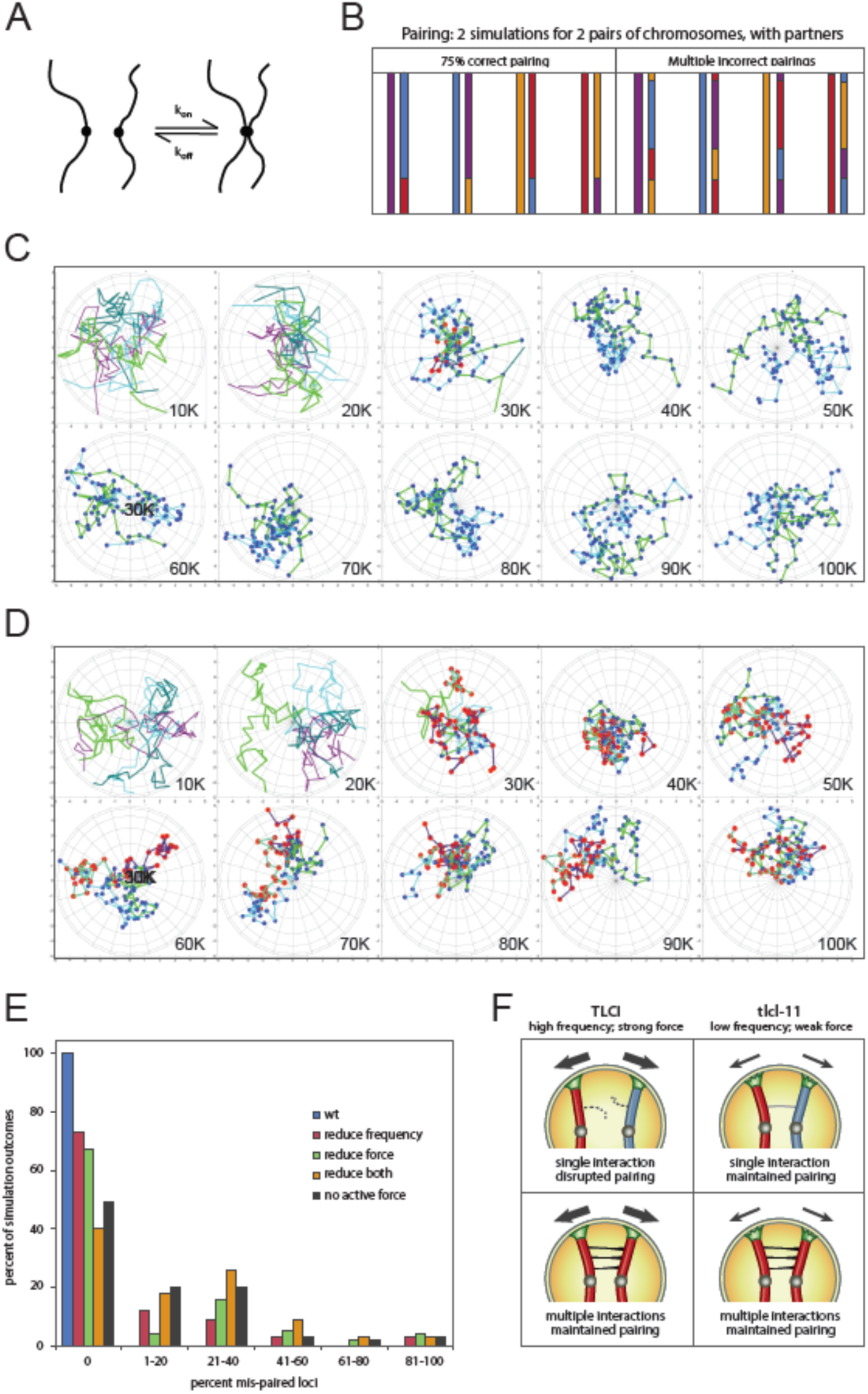
Simulation of active random forces predicts pairing errors from reduced pulling frequency. **A)** Pairing as a reversible process. Individual loci can pair when they randomly move within some capture radius. The probability of such a pairing event depends on the random motion of the chromosomes (Marshall and Fung, 2016). Once paired, any locus has a probability p(unpair) of becoming unpaired at any given time in the simulation. Correctly paired loci would have a small value of p(unpair) and incorrect loci a high value. **B)** Pairing diagrams for simulations with two pairs of chromosomes. Blue and purple lines represent one pair of homologs, while orange and red represent a second pair of homologs. Correct pairing is blue pairing to purple and orange pairing to red. Diagrams plot final results as four parallel lines, where each pair of lines represents one of the four chromosomes on the left along with a plot of its pairing partner on the right. The left panel depicts correct pairing for approximately ¾ the length of all four chromosomes, with mispairing at one end. The right panel shows multiple correct and incorrect pairings for all chromosomes. **C)** Example of simulation with 100% velocity and normal pulling frequency. Dark green and bright green represent one homolog pair and purple and blue represent a competing homolog pair. Red dot indicates a mis-paired locus and a blue dot indicates a correctly paired locus. In this WT simulation, both homolog pairs eventually become correctly associated. Examples taken at every 10,000 timepoints (10K) out of 100K. **Movies in Figure S6A-B. D)** Example of one simulation run with 100% velocity but no pulling. Without any pulling forces, example runs will show incorrect association at the end of the run. **E)** Simulating five scenarios of the effect of alteration in magnitude and frequency of telomere-led active forces. WT denotes an active telomere force equal to four times the thermal random force and a pull frequency of 1, meaning that the pulling force is constantly being exerted. “reduce frequency” keeps the high pulling force but reduces the pulling frequency to 25%. “reduce force” keeps a pulling frequency of 100% but reduced the pulling force to 75% of its full value. “reduce both” means that pulling frequency is 25% and pulling force is 75% of maximal. “no active force” means that no additional force is exerted on the telomeres other than thermal forces. The graph plots the fraction, out of 100 independent simulation runs, in which the final paired state has a given fraction of loci incorrectly paired. The first bin contains only those outcomes in which pairing was perfect, i.e. no mistakes. All other bins contain the range of mis-pairing fractions indicated. **F)** Proposed model for tlc1-11 phenotype in which reduced force and frequency of pulling motion prevents dissociation of incorrectly paired regions.

## Discussion

### Mutations reducing telomerase engagement at telomeres modulate cytoskeletal forces during meiosis

Here we have shown that in *Saccharomyces cerevisiae*, reducing telomerase levels at telomeres modestly reduces spore viability and alters meiotic recombination frequencies. Roles for telomerase components in anchoring telomeres to the nuclear envelope (NE), have been documented in mitotic cells. Our findings are consistent with the telomerase holoenzyme being important in meiosis as well, specifically for efficient engagement with the cytoskeleton via telomeric nuclear envelope attachment. We propose that telomerase-mediated engagement helps mediate proper transmission of cytoskeletal forces to chromosomes during meiosis. We report that a hypomorphic telomerase core component mutation (*tlc1-11*) that partially reduces telomerase at telomeres dramatically decreases the frequency of high velocity telomeric “pulls”, resulting in earlier homolog synapsis and increased recombination for the majority of the cells. The lack of a delay in attaining overall pairing suggests that the primary function of the pulling motion increase active diffusion to drive homologous together but instead is consistent with some of the other proposed models for this rapid telomere motion (Figure 1B)

In most organisms, the cytoskeleton is attached to chromosome ends through the nuclear membrane via the SUN/KASH domain proteins that connect to proteins that bind to the telomere. In budding yeast, although it was long known that Ndj1 is important for tethering the telomere to the nuclear membrane in meiosis, there has been a question of whether other telomeric components are required. In mouse, MAJIN (Membrane-Anchored Junction Protein) was found to link telomeric DNA to the NE (49). We found that telomerase is important for full conduction of cytoskeletal forces during prophase I, revealing an important mechanical role for telomerase in meiosis. The reduced telomerase function in *tlc1-11* alters both the velocity and frequency of the pulling component of telomere-led motions, which we propose causes the observed earlier initiation of synapsis, subpopulation of cells that are unable to exit synapsis, increased crossing over, and lower spore viability. In contrast, mutating the Esc1-Sir4 subtelomeric attachment did not reduce spore viability or increase overall recombination, suggesting that this subtelomeric anchoring mechanism is not critical for meiosis. One reason the Esc1-Sir4 attachment might be superfluous in meiosis is the presence of Ndj1. Ndj1 may provide greater strength to the telomerase dependent attachment in order to withstand the cytoskeletal forces experienced in meiosis but not in mitosis.

### Roles of Active Motion in Ensuring Fidelity of Homologous Pairing

The importance of chromosome motion for homolog pairing was first proposed after the discovery that active chromosome motion is driven by cytoskeletal forces during prophase I in *S. pombe* (*50*). Since then, that active motion speeds up the search for homologous sequences has been further established both experimentally (15, 29, 51) and theoretically (18). While active motion may facilitate homologs locating each other within the nucleus, at least three lines of evidence argue that this is not the only essential role for all components of the telomere-led forces. First, although pairing is delayed in the *ndj1*1 mutant, which has defective tethering to the NE, the extent of pairing of individual loci eventually reaches near-WT levels (28). Second, Chacon et al, 2016 (17) showed that transiently halting active chromosome motion in fission yeast results in excessively stable associations forming at loci that were already paired when the motion was halted, thus indicating a second role for the motion that acts after initial pairing. Third, our study shows that specific perturbation of chromosome pulls leads to earlier synapsis even though timing and extent of pairing occur normally. Our results provide evidence that telomerase-mediated, telomere-led motion normally acts to delay a step occurring after initial pairing and before full synapsis. We propose that this additional step is the removal of incorrect associations and/or the correction of non-terminal synapsis which are transient small synaptic associations that occur in the progression towards full synapsis (40)

### Model for removal of incorrect associations without invoking a molecular sensor

Sato et al (2009) (16) proposed that active forces are used to sense tension between associated pairing centers. Their model uses tension to probe the correctness of association in order to generate a global signaling trigger for initiation of synapsis. An alternative model is that active forces simply act to pull apart incorrectly associated regions, but are not strong enough to pull apart long stretches of correctly paired regions, such that active force favors the persistence of correct, but not incorrect, associations (Figure 7F). This simpler model does not posit any signals or tension sensors, but simply suggests that the active forces would reduce the extent of incorrect pairing relative to correct pairing. Computational simulations of this model indicate that random forces can provide a high level of selectivity in favoring correct over incorrect associations (48).

In summary, manipulation of nuclear attachment at the telomeres suggests alternative roles for chromosome motion during meiosis. Further future investigations directly examining unpairing and non-terminal synapsis will could reveal the relative impact that chromosome motion contributes to these processes.

## Materials and Methods

### Strain Construction

All yeast strains are derivatives of the SK1, BR1919-8B, S96/YJM789 hybrid or S288C backgrounds as detailed in Table S1. Complete disruption of ORFs was carried out by PCR-mediated gene disruption. For *tlc1-101* and *tlc1-11*, both integrated and plasmid-containing strains were made. To integrate the *tlc1* mutant alleles, loop-in, loop-out of a mutation-containing plasmid was followed by PCR verification to replace TLC1. Alternatively, a plasmid containing the specific mutant allele was transformed into *tlc1Δ.* For *tlc1* alleles, haploid strains were streaked serially 6 times to ensure complete penetrance of mutant telomere phenotypes (2 days of growth on solid media at 30° C represents approximately 20 generations). Haploid strains with pre-elongated telomeres were made through expression of a CDC13-Est2 fusion under the CDC13 promoter (24) in EST2 or *est2Δ* backgrounds, followed by serial streaks and southern blotting with a telomeric probe to monitor telomere length. Haploid strains of similar telomere length were mated just prior to meiosis analysis. To create diploid strains for subsequent sequencing, imaging and meiotic progression assays, otherwise isogenic MAT**a** and MATα parents with telomeres of similar telomere length were mated. For recombination mapping, S96/YJM789 hybrid diploids were sporulated. Spore viability tests were carried out in BR1919-8B and S96/YJM789. The BR1919-8B background was used for genetic recombination, chromosome spreads, synapsis measurement and movement assays. Synchronous meiotic progression assays, FACS analysis of replication onset, pairing, DSB formation and asci formation were carried out in SK1. Strains are listed in Table S1. Detailed strain genotypes as well as plasmid and oligo sequences are available upon request.

### Southern Blotting Analysis of Telomere Length

Genomic DNA was prepared from haploid cells from serial streaks on solid media after the indicated number of passages. Genomic DNA was then digested with XhoI and run on 0.8% agarose gels. DNA was transferred from the gels to Hybond N+ membranes and probed with γ32P end-labeled WT telomeric repeat oligonucleotide (TGTGGTGTGTGGGTGTGGTGT) and visualized using a phosphoimager.

### Meiotic Time Courses

To induce synchronous meiosis, overnight cultures from zygote colonies of freshly mated parent strains were grown in YPADU (for 1919) or YPD (SK1) with Adenine, .01%. After 24 hours, cultures were diluted to A600 = 0.25 in 10 mls of fresh prewarmed (30°) BYTA (1% yeast extract 1% potassium acetate, 2% Bactopeptone, 50mM potassium phthalate, .01% Adenine), shaken at 250rpm, 30°C for 14.5 hours (for A600 = 1.8-1.9), washed once in warm water and the pellet resuspended in 10 mls 0.3% SPM at 30C° at A600 = 1.8 to start meiosis. Samples were collected and fixed in 70% ethanol for time 0 and every 15 minutes, for 2 hours.

### Chromosome Motion Assay

Cell cultures were grown, induced, mounted, and then imaged on the OMX microscope and analyzed as described in Pollard and Fung (2017) and Pollard et al. (2023) (40, 52). Imaging of live cells was performed between 14 and 24 hours after induction, generally using a 4 to 5 µm z-stack with sections at 0.2 µm intervals, exposures of 5ms, and time points every 400 ms. Bleaching of the GFP fluorophore under these conditions usually limited time course length to a maximum of 10 minutes. Image data was visualized and processed by deconvolution and denoising with algorithms embedded in the Priism imaging analysis platform. Chromosome contours were semi-automatically generated using MATLAB routines and algorithms. Each contour was generated via a process in which the position of a synaptonemal complex at each time point was automatically digitized, its movement then tracked through space and time, and kinetic parameters of the motion then quantified. For motion metrics of SC ends, an approximate equal number of minutes were evaluated for each comparison strains using 4 cells for *zip3* and 11 cells for *tlc1-11 zip3*.

### Meiotic chromosome spreads

For pachytene spreads, 2 ml BR1919-8B strains were grown in YPD for 20-24 hours and diluted 1:3 fold for grown for 7 hours before transferring cell pellet from 1 ml of culture into 10 ml of 1% potassium acetate to induce sporulation at 30°C at 250 rpm. Cultures were spun down after 17, 18.5 and 20 hours of sporulation and resuspended in 2% potassium acetate 1M sorbitol with 10 ul 1M DTT and 10mM zymolyase 100T and incubated at 30°C for 26 minutes. The spheroplasted cells were gently pelleted and resuspended in 200 mM MES + 1M sorbitol pH 6.4 and repelleted. After supernatant removal, the pellet was resuspended with 200 ul 200 mM MES followed immediately with 400-500 ul of 37% formaldehyde. The cell suspension was divided between 3 slides. A 22x50 coverslip was placed over each slide after 10 minutes. After 30-45 minutes, the slides were washed with 4% Photoflow (Kodak, Rochester, NY) and dried. Slides were stained with primary antibodies guinea pig anti-Rap1 (1:100) and rabbit anti-Zip1 (1:50) diluted in 20% fetal bovine serum (FBS) in 1X PBS for two hours. After two 10-minute PBS washes, 200 ul of a dilution of 1:200 of anti-guinea pig AlexaFluor 546 and anti-rabbit AlexaFluor 488 secondary antibodies (Invitrogen) in 20% FBS in PBS were added to the slides and incubated for two hours. After two PBS washes, slides were dried briefly and mounted in 1.5 μg/ml DAPI, 1 mg/ml p-phenylenediamine in PBS-buffered glycerol. Images were collected on a DeltaVision microscope (GE Healthcare, Marlborough, MA).

### Pairing Assay

The one spot-two spot assay was used to measure pairing in SK1 strains with Chromosome V (CEN) or Chromosome IV (TEL) marked with 256 repeats of the LacO binding site. LacI-GFP was expressed from the *CYC1* promoter from the URA3 locus, ura3-1::p*CYC1*-GFPlacI::URA3. To induce synchronous meiosis, diploid strains were pre-inoculated at OD600 = 0.3 in BYTA medium and grown as described above. Samples were collected hourly for 8 hours. At each time point, cells were harvested, washed, pelleted and resuspended in .5 ml of 4% paraformaldehyde, 3.4% sucrose, and held at room temperature for 10 min. Cells were washed once in .5 ml of 0.1 M potassium phosphate, 1.2 M sorbitol buffer and resuspended in the same buffer. Cells were sonicated before microscopy, and spun onto concanavaline-A coated slides before being mounted with Vectashield plus DAPI. Images were collected on a Deltavision microscope using a 60x objective and images were analysed using Fiji and CellProfiler. The number of GFP foci/nucleus was averaged from ∼1000 nuclei/timepoint.

### Synapsis Progression

Cell cultures were grown, induced, mounted, and then imaged on the OMX microscope as described in Pollard and Fung (2017) (52). Imaging of live cells was performed beginning after 5.5 hours post induction, in 10µm z-stacks with sections at 0.2 µm intervals with exposures of 5ms, and time interval every 30 minutes. Time courses were routinely terminated after 150 hours of imaging (time points were generally acquired with stepped-down frequency at later times: 1, 2 or 4 hour intervals). Image data was visualize, denoised and enhanced using Priism software. Individual cells were both manually and algorithmically tracked and scored for progression past: 1) the start of Zip1 expression, 2) the onset of synapsis, 3) the termination of synapsis, and 4) the formation of spores and/or the appearance of auto-fluorescence in spores. Scripts written using MATLAB software (MathWorks Inc., Natick, MA) were utilized to analyze population progression data statistically.

### Whole-Genome Microarray and Next Generation Sequencing Recombination Mapping

DNA was prepared for Illumina sequencing using a NextFlex kit (BIOO Scientific, Austin TX) with Illumina-compatible indices or as described (45) with 4-base or 8-base inline barcodes. Read alignment, genotyping and recombination mapping were performed using the ReCombine package (45). While running CrossOver.py, the input values for ’closeCOs’, ’closeNcoSame’ and ’closeNCODiff’ were all set to 0. Insertions and deletions were removed from the set of genotyped markers. Recombination events within 5kb of each other were then merged into single events and categorized into seven types as described (53). Gamma distributions and CoC were calculated as described (46). Standard error for CoC was calculated from the average of the 25 kb bins used to calculate interference. S96 microarrays (Affymetrix, Santa Clara, CA) were prepared and analyzed with Allelescan software.

### Microarray Detection of ssDNA

A total of 1.5 mg each of 0 hr and 3 or 5 hr ssDNA samples were labeled with Cy3-dUTP or Cy5-dUTP (GE Healthcare) by random priming without denaturation with 4 mg random nonamer oligo (Integrated DNA Technologies, Skoie IL) and 10 units of Klenow (New England Biolabs, Ipswich, MA). Unincorporated dye was removed with Microcon columns (30 kDa MW cutoff; Millipore, Burlington, MA), and samples were cohybridized to custom Agilent arrays in accordance with a standard protocol. For each set of experiments, a dye swap was performed.

### Southern Blotting Analysis for DSB Measurement

To induce synchronous meiosis, diploid strains were pre-inoculated at OD600 = 0.3 in BYTA medium (50 mM potassium phthalate, 1% yeast extract, 2% bactotryptone, 1% potassium acetate), grown for 16 hr at 30 C, washed twice, and resuspended at OD600 = 1.9 in SPO medium (0.3% potassium acetate). Southern analysis was performed by separating DNA fragments in 1% agarose/1X TBE gels using pulse-field electrophoresis with a 5-45 s ramp and blotted onto Hybond-XL membranes (GE Healthcare) with alkaline transfer.

### Quantification and Statistical Analysis

All statistical analysis was performed using R, Priism (GraphPad) or Microsoft Excel.

## Data and Software Availability

Raw sequence data from the genome-wide analysis of recombination have been deposited in the NIH Sequence Read Archive under accession number SUB4518296. Custom MATLAB and Allelescan software is available freely upon request. Recombine software is available at https://sourceforge.net/projects/recombine/files/1574 Word count for Materials and methods

## Acknowledgments

We thank Beth Rockmill for comments on the manuscript.

Support is provided by R01 GM116895 and R01 GMGM137126 to AO, TG, WM & JCF; RO1 GM026259 to DLS & EHB R01 GM111715 to AH & NJP and R01 GM075119 to SB & DC.

---

**Figure S1.**
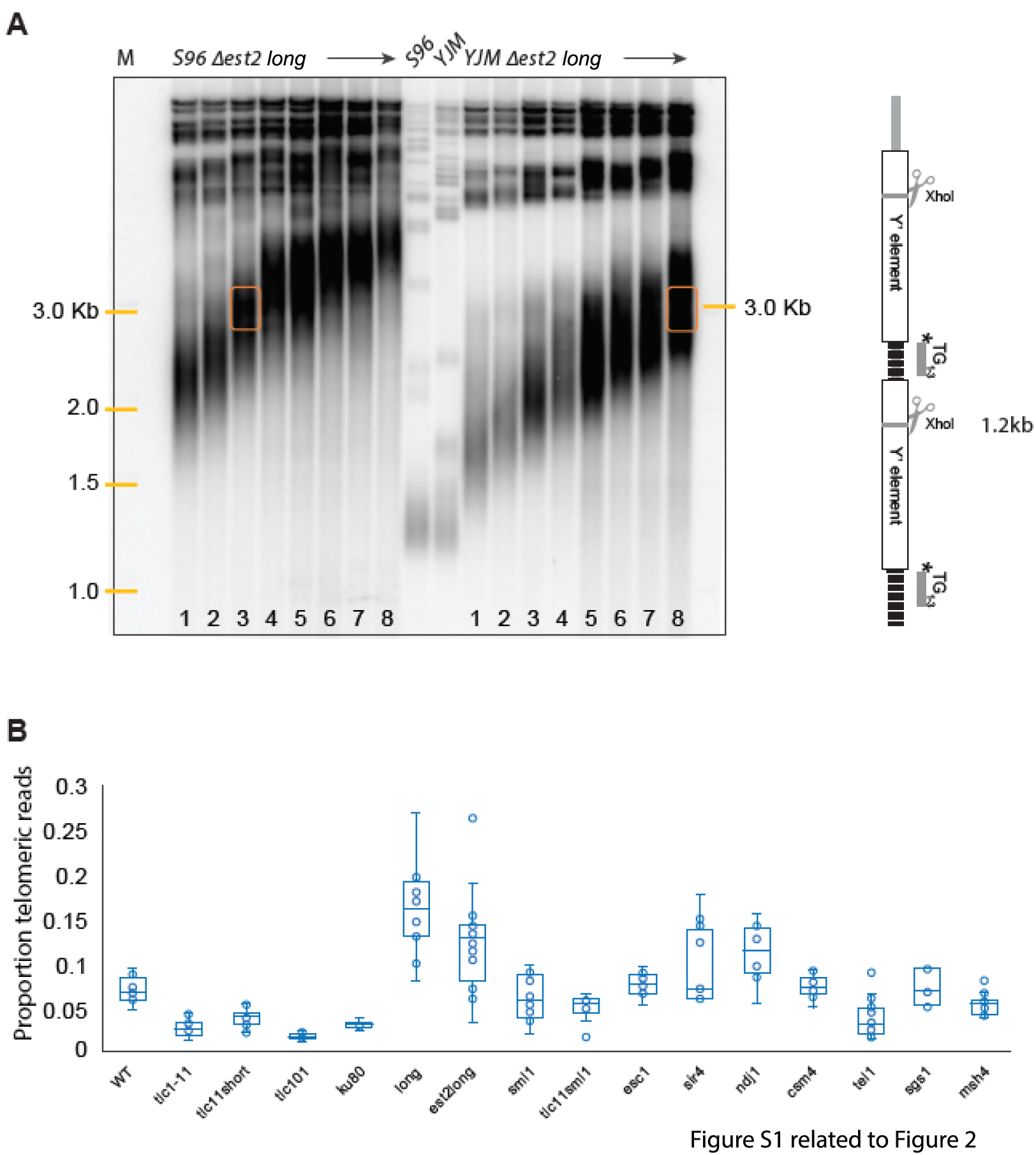
Construction of strains with long telomeres but no telomerase. **A)** *EST2* was deleted in S96 and YJM789 parents and plasmid-borne Cdc13-Est2 fusion protein was integrated at CDC13. Cells were serially passed 8 times (totaling about 160 generations) to ensure extensive lengthening of telomeres. The Cdc13-Est2 fusion protein was selected against, ensuring complete loss of telomerase in a long-telomere context. S96 MAT**a** and YJM789 MATα parents with similarly long telomeres were selected for mating and subsequent sequencing of meiotic products. Shown is Southern blot analysis of terminal XhoI restriction fragments. DNA was probed with an γ^32^P-labeled 5’-(TGTGGG)_4_-3’ Y’-specific probe. The lowest species represents the DNA fragment containing the terminal telomeric repeats. The selected parents have telomeres that are about 1750 bps, or 5x WT. **B)** Swarm plot of data shown in Figure 2D.

**Figure S2.**
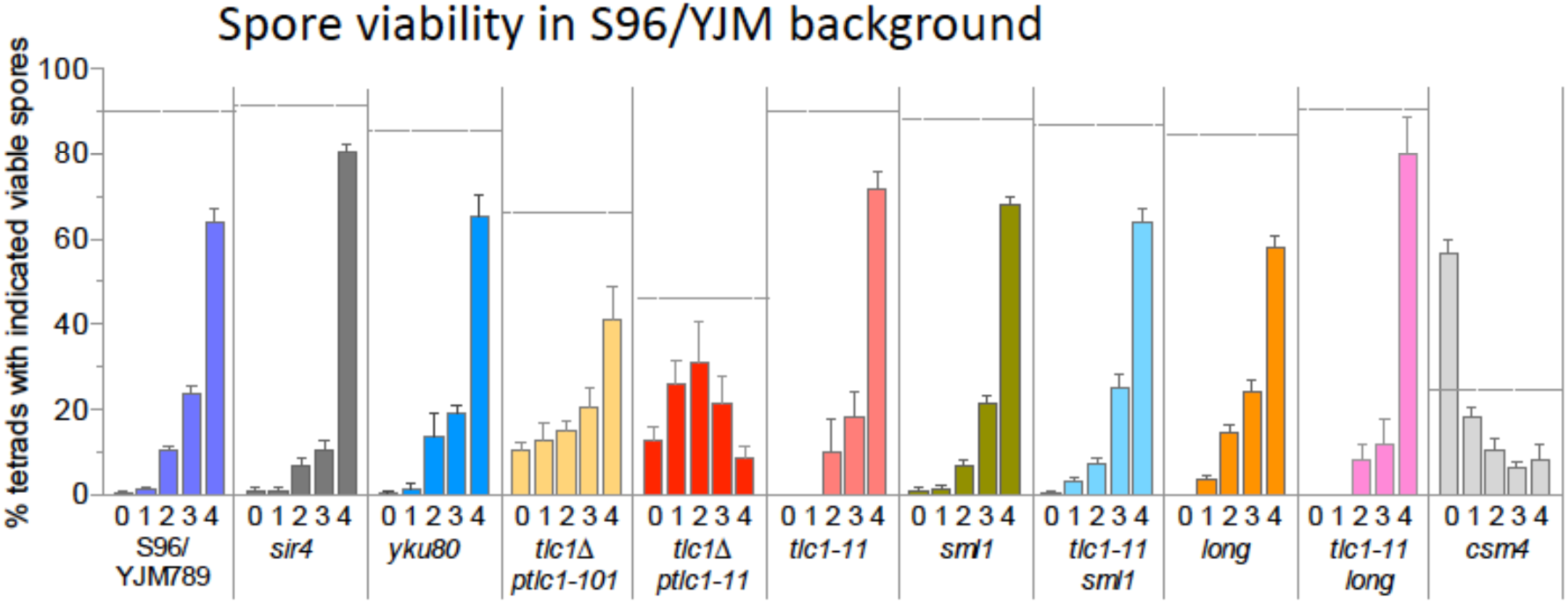
(Relates to Figure 2) Spore viability in hybrid strain background. The spore viability measurements in S96/YJM789 backgrounds. *yku80Δ, tlc1Δ ptlc1-101, tlcΔ ptlc1-11, sml1Δ, tlc1-11 sml1Δ, long, tlc1Δ long,* and *csm4Δ* mutants lead to spore inviability, whereas *sir4Δ* does not. Spore viability, % asci with 0,1,2,3 or 4 viable spores as indicated for each genotype below the histograms. Horizontal dotted line indicates total spore viability for each strain. (See Table S1 for more mutant spore viability data)

**Figure S3.**
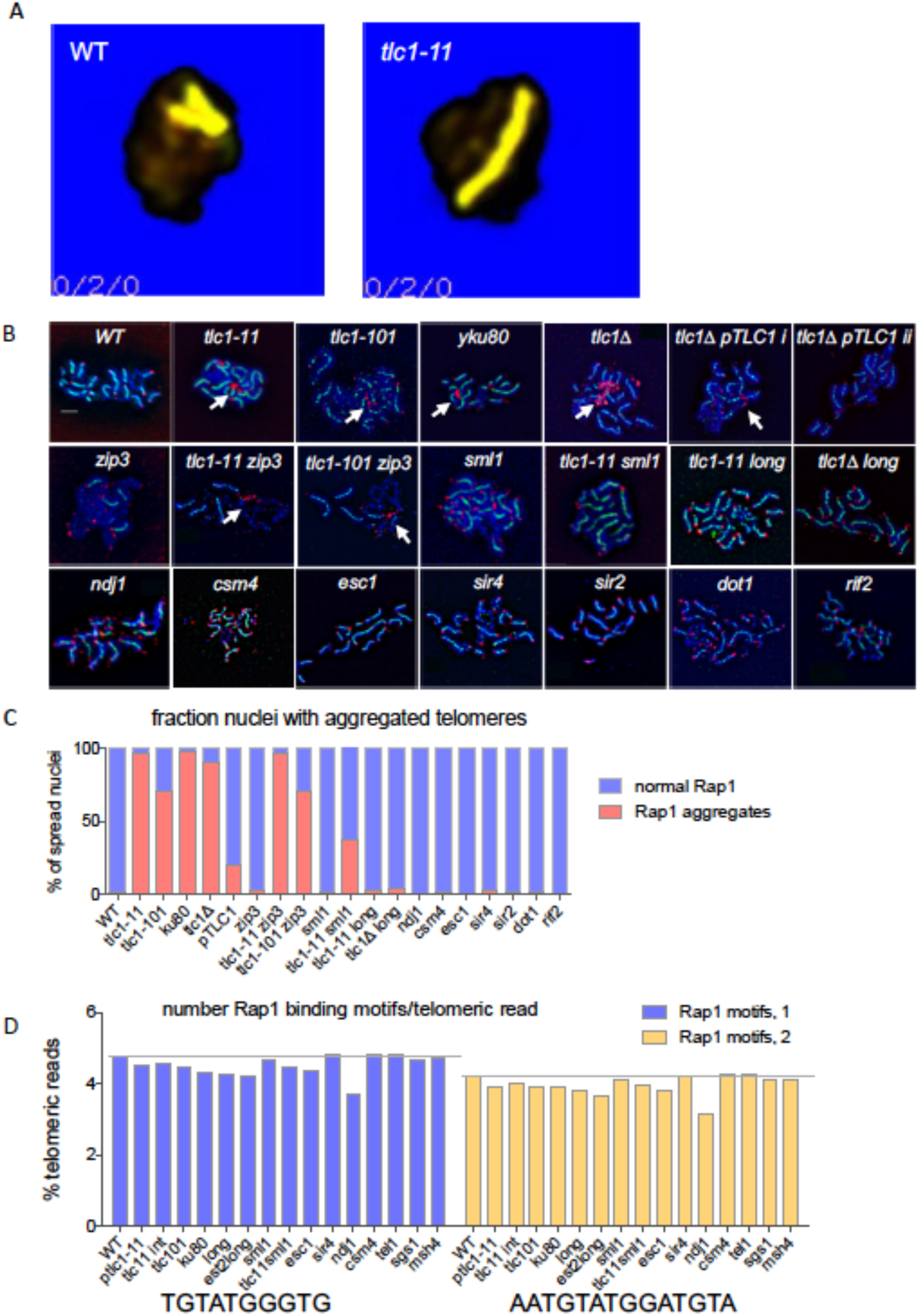
Meiotic SC movement and mutations affecting telomere length result in Rap1 aggregation. **A)** Meiotic SC movement. Max intensity projection of a zip3Δ nucleus with a single synapsed synapsed chromosome marked with Zip1-GFP for WT (right) and *tlc1-11*. See Supplemental Movies Figure S3A-B;. **B)** Pachytene chromosomes were spread and immunostained. Blue is DAPI staining of DNA, Green shows immunostaining of Zip1-GFP and Red represents immunostaining of Rap1. White arrows indicate representative telomeric aggregates of Rap1. Rap1 binds site-specifically to double-stranded DNA throughout the genome, at discrete promoter sites (Huet et al., 1985). The highest concentration of Rap1 binding sites occurs in telomeric and sub-telomeric regions, although Rap1 is not thought to anchor telomeres to the nuclear envelope. In WT pachytene nuclei, Rap1 appears as discrete spots at the end of each chromosome (Figure 4H). In *tlc1-11*, Rap1 staining at telomeres is faint, consistent with the shorter telomere length in the mutant. Interestingly, in most *tlc1-11* nuclei the Rap1 protein largely appears in large aggregates (96%, Figure 4H). Rap1 aggregation occurs in *tlc1-101* (70%) and y*ku80*Δ (97%), but is very rare in WT (2%) (Figure 4H). Virtually no Rap1 aggregation is observed in deletion mutants, Sir4 or Esc1, Csm4, Ndj1, or in mutants of the sub-telomere maintenance proteins Sir2, Dot1 and the telomere length regulation protein Rif2. **C)** Quantification of meiotic Rap1 aggregates. Percentage of pachytene spread nuclei with RAP1 aggregates present indicated in red and fraction without aggregates are indicated in blue. **D)** Analysis for changes in the Rap1 binding motif in mutant strains Two Rap1 binding motifs whose sequence is shown below the graph were used to determine the percentage of reads containing these motifs in WT and mutants. Motif 1 is from the JASPAR database of transcription factor binding sites and Motif 2 is from Lascaris et al. (1999) that analyzed Rap1 binding.

**Figure S4.**
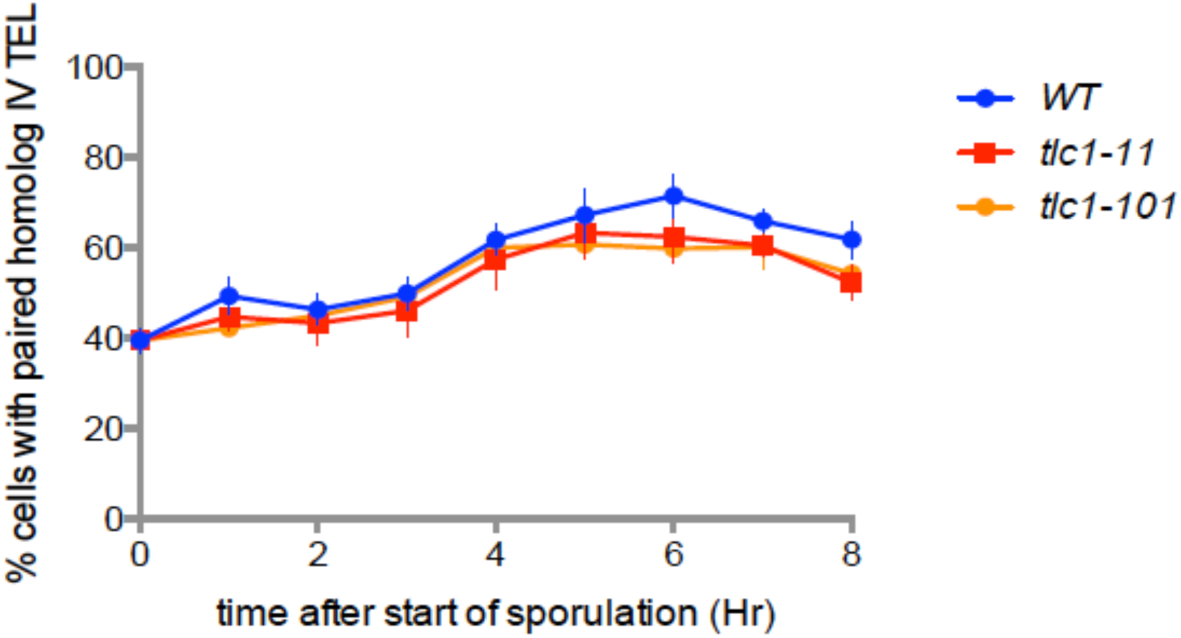
Pairing of homologs, marked at telomeres. Pairing of homolog IV TEL for WT, *tlc1-11 and tlc1-101*. Cells were gathered and fixed every hour after start of meiosis in cells with LacO repeats inserted near the telomere of each homolog. LacO foci detected with LacI-GFP are considered paired (1 spot observed) or unpaired (two GFP-lacI spots observed) n=1000 per time point.

**Figure S5.**
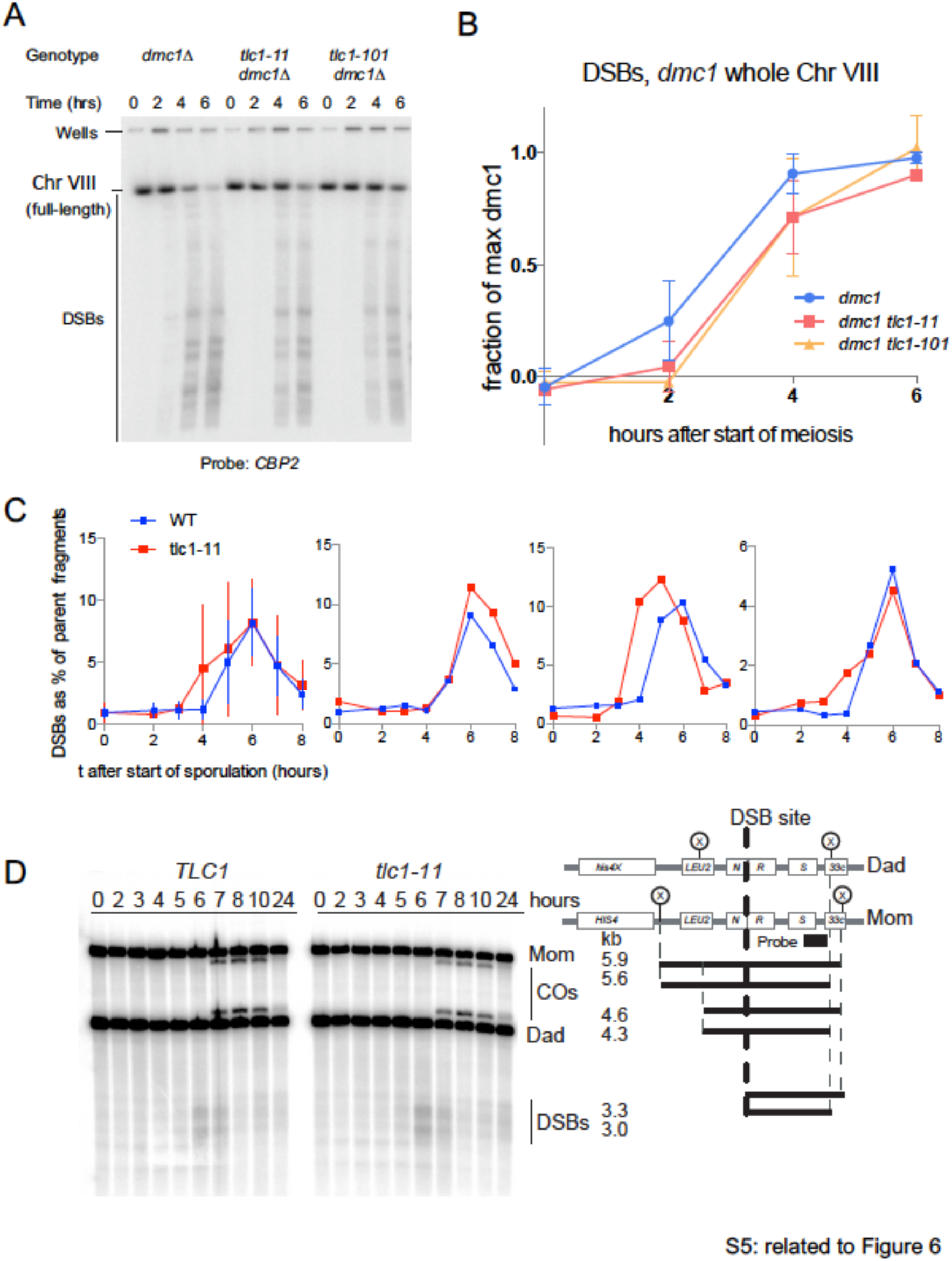
Meiotic DSBs, on chromosome 8 and at *HIS4-LEU2* hotspot. **A)** Southern blot comparing DSB number, over time, in *dmc1, tlc1-11 dmc1*, *tlc1-101 dmc1.* A *dmc1*Δ background was used to limit DSB processing. Pulse field gel electrophoresis of DNA from samples collected every 2 hours was probed with CBP2. **B)** The timing and number of DSBs on Chromosome VIII in *tlc1 dmc1* mutants is expressed as the fraction of *dmc1* maximum and is not significantly different from WT. **C)** Replicate time courses, showing quantification of DSBs at HIS4-LEU2 break site of Chromosome III. An average of three time courses is shown in left-most graph, with graphs of each of the three individual time courses shown to the right. **D)** One-dimensional gel analysis of a blot from time course 1 (TC1) depicting meiotic recombination at the *HIS4::LEU2/his4-X::LEU2-URA3* hotspot in *TLC1* and *tlc1-11* strains. Time course samples were treated with psoralen. DNA isolated at various timepoints was digested with XhoI and separated on a 6% agarose gel. Southern blots were hybridized with probe A (3’ end of *STE50* gene open reading frame). Schematic identification of break fragments is as described by Oh et al. (2009).

**Figure S6.**
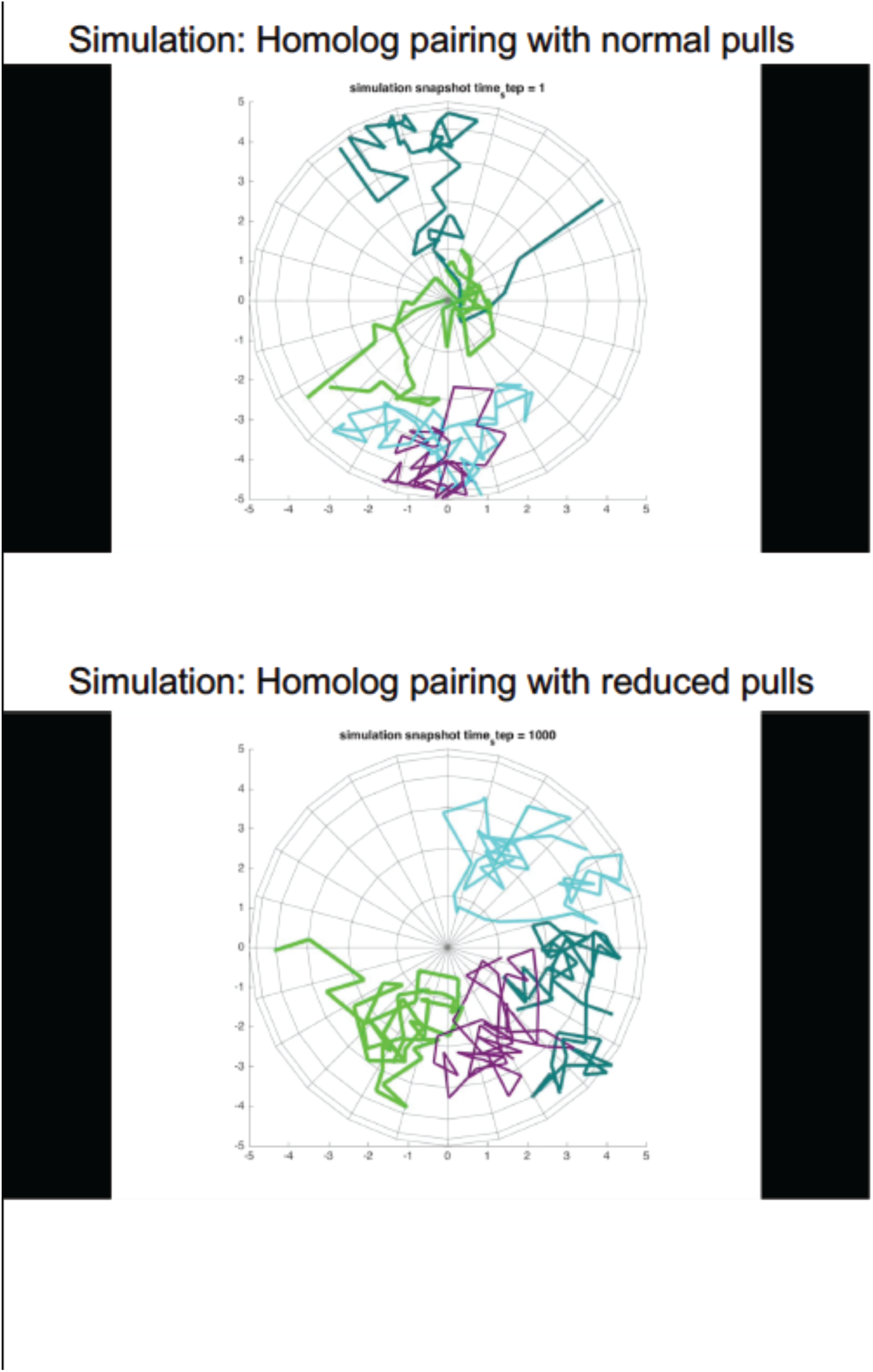
Simulation movies with two pairs of chromosomes. (one pair represents true homologous partners and the other set represent homeologous partners) The movies depict simulation of homolog pairing with normal pulls **(A)** and homolog pairing without pulling **(B)**. **Movie: A:** Example of simulation r with 100% velocity and normal pulling frequency. Dark green and bright green represent one homolog pair and purple and blue represent a competing homolog pair. Red dot indicates a mis-paired locus and a blue dot indicates a correctly paired locus. In this WT simulation, both homolog pairs eventually become correctly associated. Examples taken at every 10,000 timepoints (10K) out of 100K. **Movie: B.** Example of one simulation run with 100% velocity but no pulling. Without any pulling forces, example runs will show incorrect association at the end of the run.

**Table S1.**
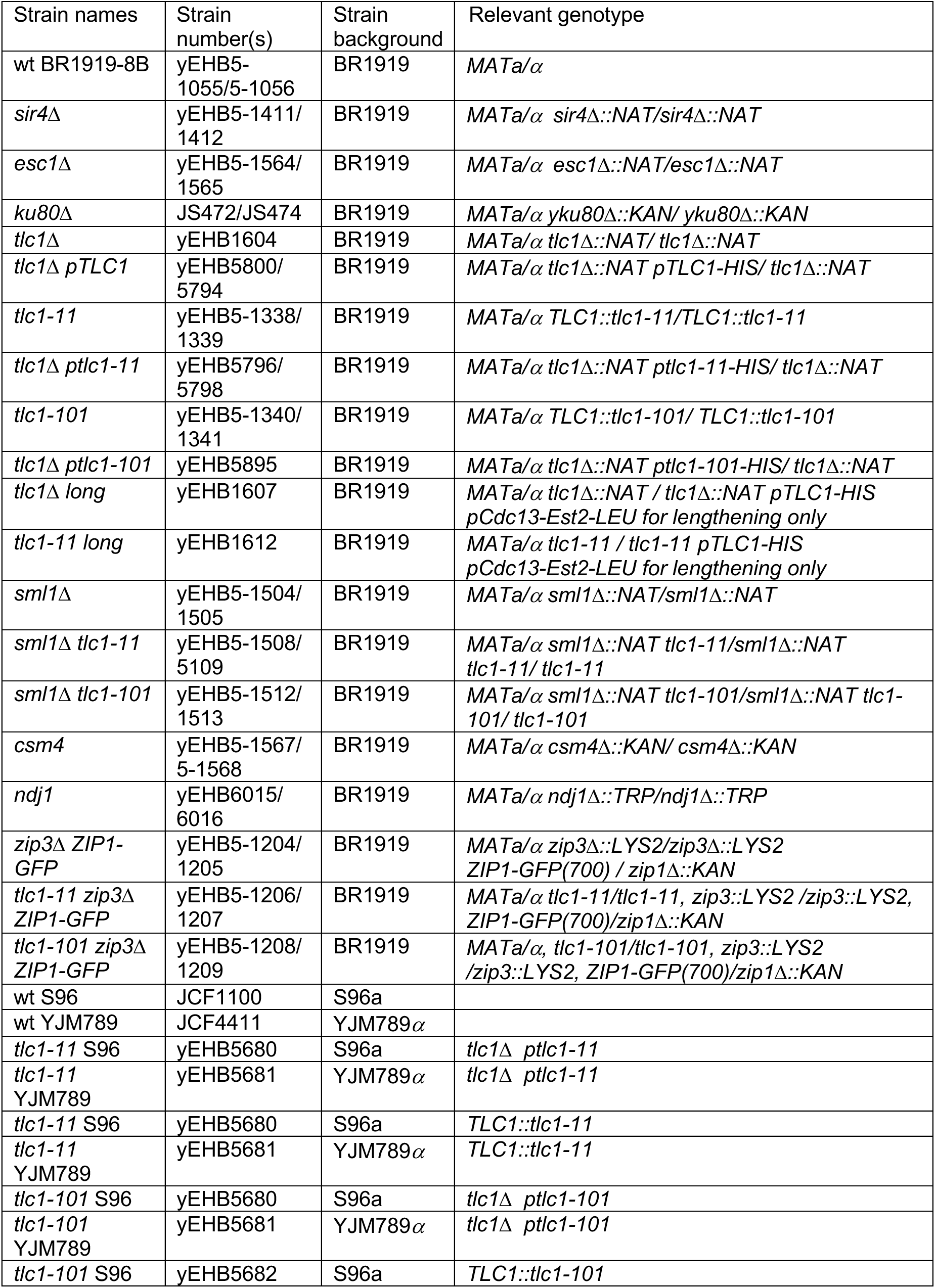
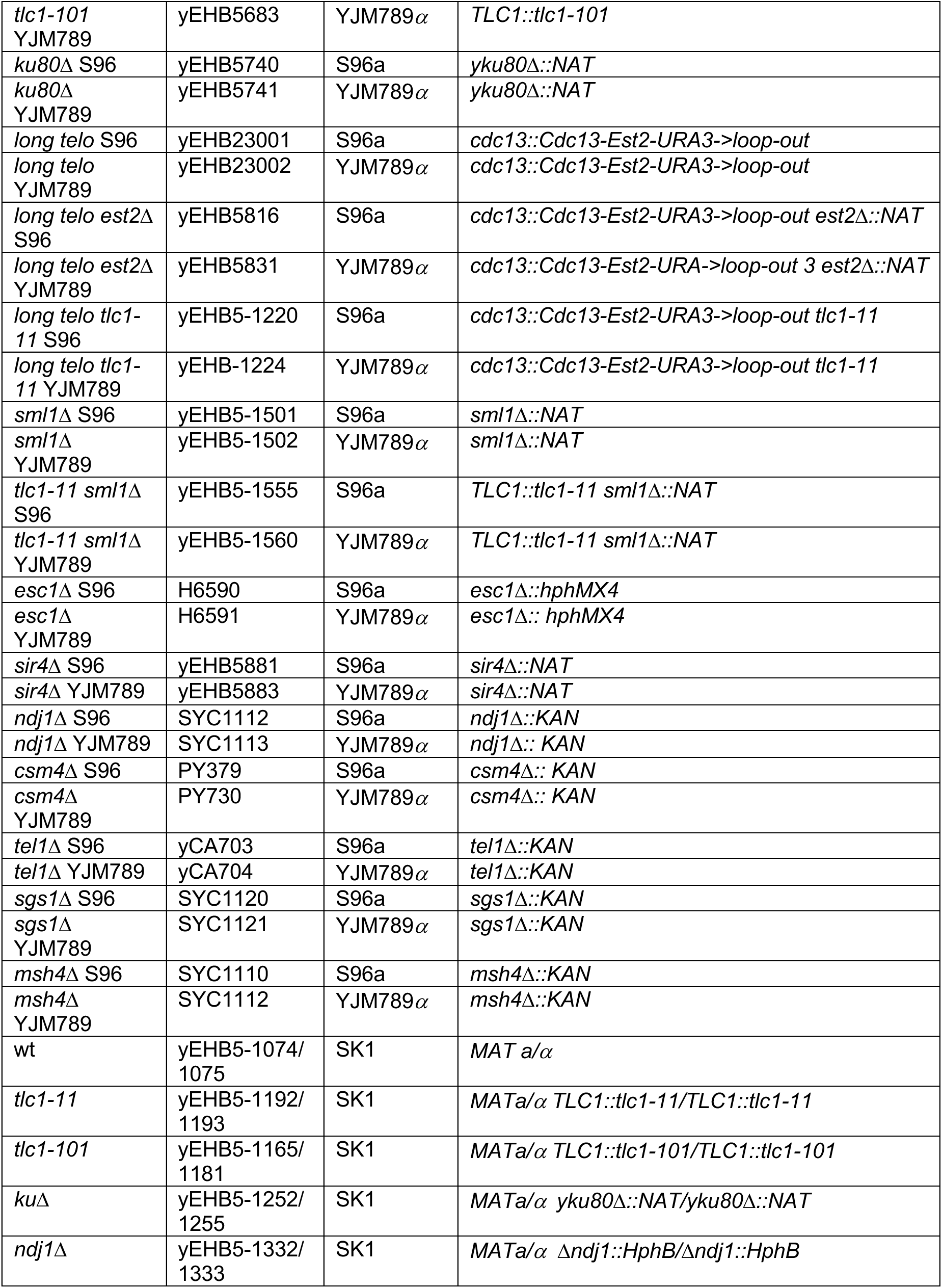
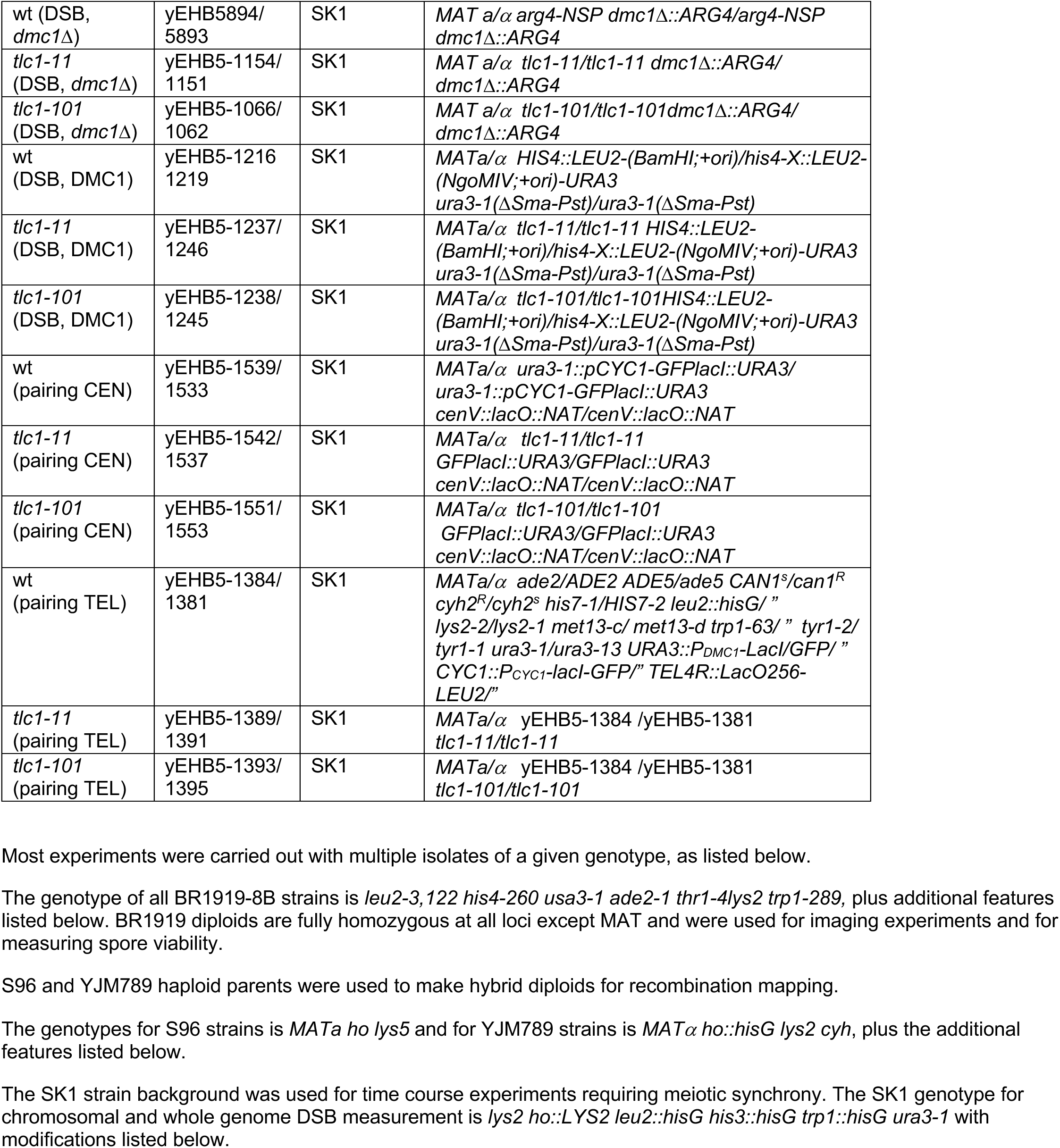
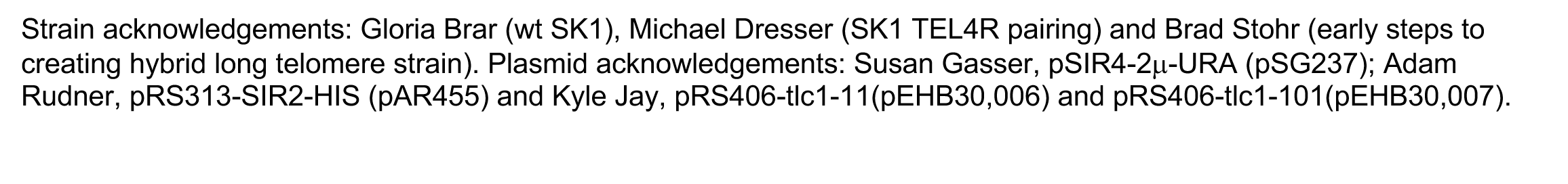
Yeast strains used in this study.

**Table S2.** Spore viability, related to Figure 2 and S3.

| % Viable spores / tetrad | 0 | 1 | 2 | 3 | 4 | Total Viability | Notes |
| --- | --- | --- | --- | --- | --- | --- | --- |
| *1919* | 1 | 0 | 2.5 | 12.5 | 84 | 94.4 | WT used for microscopy |
| *S96xYJM789 (SxY)* | 1 | 0 | 10 | 15 | 74 | 90 | WT used for sequencing and CO analysis |
| *ptlc1-11 (SxY)* | 13.5 | 17.5 | 29 | 33 | 7 | 49 | Plasmid-*tlc1-11* covering *tlc1Δ* |
| *ptlc1-11 (1919)* | 23.7 | 14.6 | 24.1 | 18.3 | 18.3 | 48 | Plasmid-*tlc1-11* covering *tlc1Δ* |
| *tlc1-11 (1919)* | 0.4 | 2 | 13.8 | 25.6 | 58.5 | 76.8 | Mutation integrated at TLC1 |
| *ptlc1-101 (SxY)* | 11.5 | 12 | 15.5 | 20.5 | 40 | 67 | Plasmid-*tlc1-101* covering *tlc1Δ* |
| *tlc1-101 (1919)* | 0 | 0.75 | 5.6 | 23 | 70.5 | 83 | Mutation integrated at TLC1 |
| *yKu80Δ (1919)* | 0 | 0 | 5.9 | 12.8 | 81.9 | 94 | Deletion, marked with KAN |
| *yKu80Δ (SxY)* | 0 | 1 | 15 | 19.5 | 64.5 | 87 | Deletion, marked with KAN |
| *long telo (1919)* | 1 | 2 | 5 | 30 | 62 | 87.5 | pCdc13-Est2 |
| *long telo (SxY)* | 0.5 | 3.5 | 14.5 | 24 | 58 | 84.2 | pCdc13-Est2 |
| *long telo Δest2 (SxY)* | 0.4 | 3.6 | 18.8 | 19.6 | 57.6 | 82 | pCdc13-Est2 |
| *sir4Δ (1919)* | 0 | 0 | 2 | 14 | 84 | 95 |  |
| *sir4Δ (SxY)* | 1 | 1 | 7 | 10.5 | 80.5 | 92 |  |
| *esc1Δ (1919)* | 0 | 0.4 | 4.5 | 11.3 | 83.6 | 94.5 |  |
| *zip3Δ zip1-GFP (1919)* | 1.6 | 5 | 19.1 | 20 | 54.1 | 80 | Motion background |
| *tlc1-11 zip3Δ zip1-GFP (1919)* | 8.3 | 3.3 | 20 | 24.1 | 44.1 | 73 | Motion background, 4-spore decrease, 0-spore increase |
| *tlc1-101 zip3Δ zip1-GFP (1919)* | 9.1 | 5.8 | 19.1 | 19.1 | 46.6 | 72 | Motion background, 4-spore decrease, 0-spore increase |
| *csm4Δ (SxY)* | 56 | 18.5 | 10.5 | 6.5 | 8 | 24.35 |  |
| *csm4Δ (1919)* | 26 | 5 | 20 | 9.5 | 39.5 | 45.9 |  |

